# Maternal immune activation imprints a regulatory T cell deficiency in offspring that drives an autism-like phenotype

**DOI:** 10.1101/2025.01.06.631430

**Authors:** Pierre Ellul, Gwladys Fourcade, Vanessa Mhanna, Nicolas Coatnoan, Susana Bodula, Danielle Seilhean, Laura Mouton, Gwendolyn Marguerit, Richard Delorme, Tian Mi, Ben Youngblood, Michelle Rosenzwajg, Nicolas Tchitchek, David Klatzmann

## Abstract

Maternal immune activation (MIA) triggers an IL-17-driven autism spectrum disorder (ASD) in mouse and human offspring. While regulatory T cells (Tregs) regulate Th17 cells, their involvement in MIA and ASD pathogenesis is unknown. At the maternal level, we show that Treg stimulation suppresses interleukin-17 (IL-17) production and prevents ASD-like behaviors in offspring. At the offspring level, we show that MIA imprints a systemic and brain Treg deficiency, as evidenced by alterations in the methylome, transcriptome, and functional assays. This deficiency promotes brain inflammation, characterized by infiltration of IL-17–producing cells and neutrophils into the meninges and alterations in cortical brain structure. Stimulation of offspring Tregs with interleukin-2 reversed brain inflammation and cured established ASD-like behaviors. Thus, MIA-induced ASD is a neuroimmune disorder that can be reversed by immunomodulation.

**One sentence abstract:** Maternal immune activation during pregnancy imprints a Treg deficiency in offspring that perpetuates brain inflammation and an autism-like phenotype, which can be reversed by Treg stimulation.

## Main Text

The etiopathogenesis of autism spectrum disorder (ASD) depends on a complex interplay between genetic predisposition and environmental risk factors (*1*). Substantial epidemiologic evidence indicates that maternal immune activation (MIA) during pregnancy (e.g., from infections or autoimmune disease flares) is associated with an increased risk of ASD in the offspring (*2*). For example, children whose mother was infected with rubella during pregnancy have a 10% risk of developing ASD (*3*, *4*), while those whose mother has an autoimmune disease have a twofold increased risk (*5*).

The causal relationship between MIA and ASD is supported by experimental models. Poly(I:C) injection into pregnant mice, to induce MIA, triggers symptoms in the offspring (*6*) that resemble core ASD symptoms in humans (*7*), including communication deficits, repetitive behaviors, and impaired social interactions. Mechanistically, the current view is that MIA disrupts neurodevelopment by activating maternal effector T helper (Th17) cells, which in turn produce interleukin-17a (IL-17A). This pro-inflammatory cytokine crosses the placenta and binds to IL-17A receptors on fetal neurons, thereby affecting neurodevelopment (*8*). The ASD phenotype is driven by long-term activation of the IL-17A pathway in the brain, which induces disruption of synaptic connectivity (*9*, *10*).

Proinflammatory Th17 cells are controlled by regulatory T cells (Tregs), a subset of CD4+ T cells found in most tissues that plays a pivotal role in maintaining immune system homeostasis and controlling inflammation and peripheral tolerance (*11*). Brain-resident Tregs are involved in several processes, including (i) astrogliosis suppression (*12*), (ii) oligodendrocyte differentiation and myelination support (*13*), (iii) microglia polarization towards a neuroprotective phenotype, and (iv) limitation of the local inflammatory response (*14*). We have previously reported a striking decrease in Treg numbers, and an altered Treg:Th17 balance, in persons with ASD (*15*). Here we present finding from in vivo mouse studies that demonstrate that Tregs play a major role in MIA-induced ASD (_MIA_ASD) at both the maternal and offspring level.

### Stimulation of maternal Tregs prevents _MIA_ASD in offspring

As maternal Th17 cells are central to the development of _MIA_ASD in offspring (*8*), and Tregs control Th17 cells (*16*, *17*), we first investigated whether maternal Tregs could mitigate the effects of MIA in mice. To stimulate Tregs, pregnant mouse dams were injected daily with a low dose of IL-2 (IL-2_LD_) or PBS between embryonic day (E) 7.5 (E7.5) and E11.5, and then each group was injected with either poly(I:C), for MIA induction, or PBS, for a control, at E12.5 **(Fig. 1A)**; the F1 offspring are referred to herein as PBS-MIA_F1_ and IL2-MIA_F1_ for the poly(I:C)-treated groups, and as PBS-Ctrl_F1_ and IL2-Ctrl_F1_, for the control groups. In the dams, MIA via poly(I:C) injection resulted in a small but significant expansion of Tregs as compared to PBS-treated dams **(Fig. 1B**), while IL-2_LD_ administration significantly expanded Tregs, to a similar degree in PBS- or the poly(I:C)-treated dams **(Fig. 1B**). In dams with MIA, the IL-17A serum levels markedly increased in PBS-injected dams as expected; however, this increase was completely blocked in IL-2_LD_-injected dams **(Fig. 1C)**.

**Fig. 1.**
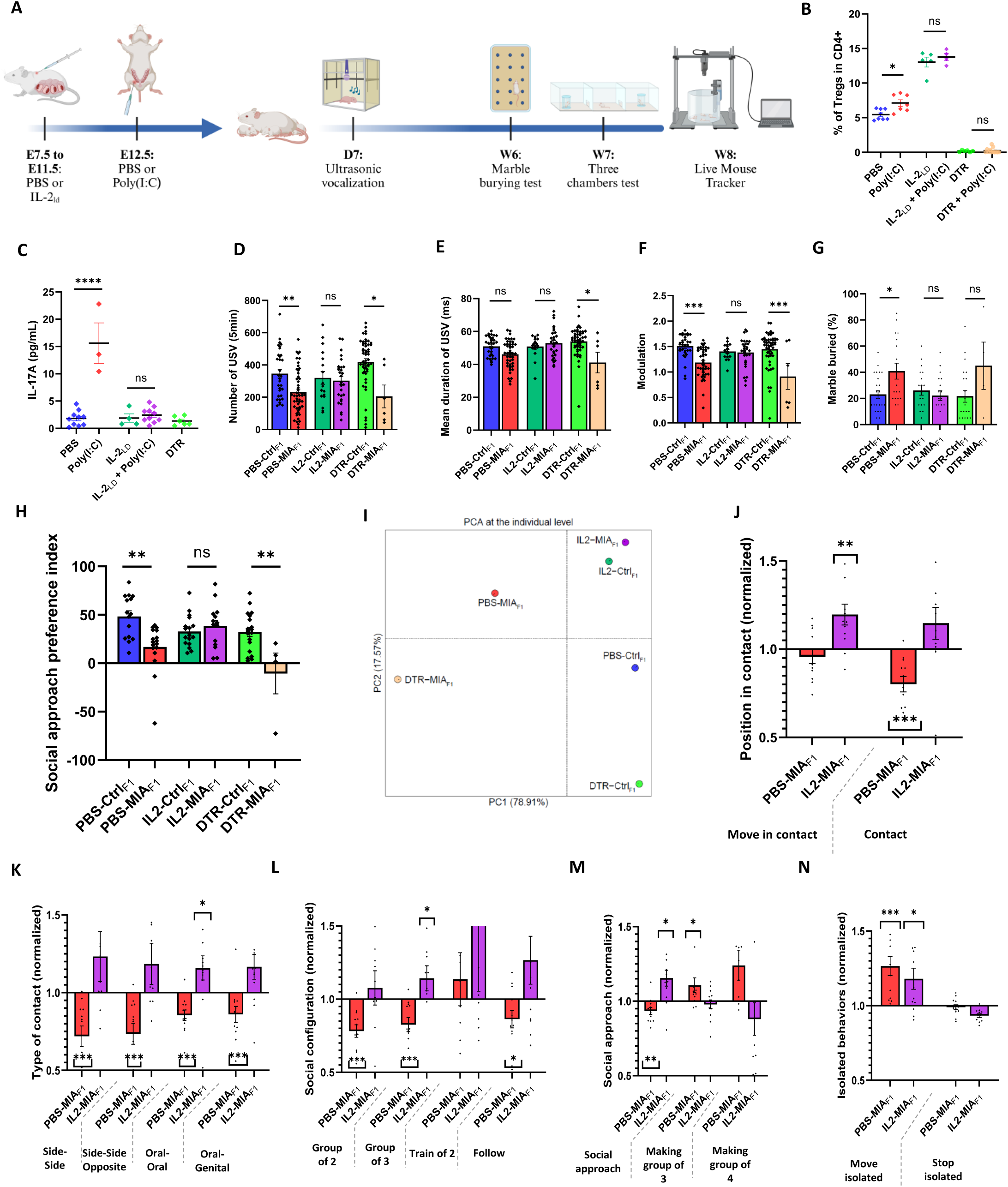
Stimulation of maternal Tregs prevents autistic phenotype in offspring. **(A)** Schematic of experimental design. Between E7.5 to E11.5, pregnant C57BL/6 mothers were subcutaneously injected with PBS or IL-2_LD_ (50,000 IU). At E12.5, pregnant mothers were intraperitoneally injected with PBS (for controls) or poly(I:C) (for MIA). F1 offspring are denoted as PBS-Ctrl_F1_, IL2-Ctrl_F1_, PBS-MIA_F1_, and IL2-MIA_F1_. At day 7 (D7), pups were separated from the mothers to record USV. Marble burying were evaluated at week 6 (W6). The F1 mice were then isolated for one week, and the three-chamber test was performed at W7. The live mouse tracker (LMT) was recorded for 14 h at W8. **(B** and **C)** Flow cytometry of maternal Tregs (B) and serum maternal IL-17A (C), at 3 h after PBS or poly(I:C) injection. **(D** to **F)** Ultrasonic vocalization (USV) characteristics of PBS-Ctrl_F1_ (*n* = 32, 5 experiments), PBS-MIA_F1_ (*n* = 49, 7 experiments), IL2-Ctrl_F1_ (*n* = 17, 4 experiments), and IL2-MIA_F1_ (*n* = 29, 7 experiments), showing number of USV recorded within 5 min (D), mean USV duration (in ms) (E), and number of modulations per USV (F). **(G)** Percentage of marbles buried in F1 mice, for PBS-Ctl_F1_ (*n* = 25, 5 experiments), PBS-MIA_F1_ (*n* = 17, 4 experiments), IL2-Ctrl_F1_ (*n* = 20, 4 experiments), and IL2-MIA_F1_ (*n* = 18, 4 experiments) **(H)** Social preference index in F1 pups, for PBS-Ctrl_F1_ (*n* = 25, 5 experiments), PBS-MIA_F1_ (*n* = 17, 4 experiments), IL2-Ctrl_F1_ (*n* = 20, 4 experiments), and IL2-MIA_F1_ (*n* = 18, 4 experiments). (**I**) PCA of the different conditions. (**J** to **N)** Live mouse tracker (LMT) behavioral profiles, for PBS-MIA_F1_ (*n* = 12) and IL2-MIA_F1_ (*n* = 12), normalized to PBS-Ctrl_F1_ or IL2-Ctrl_F1_, respectively (*n* = 12 per conditions), over 14 h of free interactions. For each mouse, we analyzed: position in contact (J), type of contact (K), social configuration (L), social approach (M), and isolated behaviors (N). Statistical analyses: one-way ANOVA with Tukey’s multiple comparisons test, except for LMT, which used Wilcoxon matched-pairs signed rank test; \**P* < 0.05, \*\**P* < 0.01, \*\*\**P* < 0.001, \*\*\*\**P* < 0.0001. Results are presented as means +/– SEM.

This stimulation of maternal Tregs had a marked effect on offspring behavior. Using a standard tool for identifying early autistic-like communication in rodent models (*18*), we next assessed their early communication skills at post-birth day 7 (D7) using maternal isolation-induced ultrasonic vocalization (USVs) **(Fig. 1, D** to **F**). Quantitatively, both the number **(Fig. 1D)** and duration **(Fig. 1E)** of USVs were reduced in PBS-MIA_F1_ pups as compared to the Ctrl_F1_ groups, but this reduction was reversed in IL2-MIA_F1_ pups **(Fig. 1, D** and **E)**. Qualitatively, USV modulation was reduced in PBS-MIA_F1_ pups but not in IL2-MIA_F1_ pups, showing that this offspring behavior was prevented by IL-2_LD_ treatment prior to MIA induction in dams **(Fig. 1F)**. Likewise, (i) repetitive stereotypic behaviors of ASD symptoms, classically assessed in mice by the marble burying test (*19*), were increased in PBS-MIA_F1_ but not in IL2-MIA_F1_ **(Fig. 1G);** (ii) social behaviors, assessed using the three-chamber test (*20*), were reduced in PBS-MIA_F1_ pups but normalized in IL2-MIA_F1_ pups, i.e., by maternal IL-2_LD_ treatment **(Fig. 1H)**. Overall, maternal Treg stimulation prior to MIA prevented the onset of the three main categories of ASD symptoms in MIA_F1_ pups.

We next investigated whether Treg ablation prior to MIA would have the opposite effect as Treg stimulation. We used Foxp3-DTR (human diphtheria toxin receptor) mice to deplete Tregs (*21*) by diphtheria toxin injections at E10.5 and E11.5. Subsequent maternal Treg depletion **(Fig. 1B)** did not per se induce changes in USV (**Fig. 1D-F**), repetitive behaviors **(Fig. 1G**), or social behaviors, of Ctrl_F1_ (**Fig. 1H).** The combination of Treg depletion and poly(I:C) injection at E12.5 resulted in a 90% abortion rate, making it difficult to study offspring behavior. Nevertheless, in the few surviving offspring (herein, DTR-MIA_F1_), we observed a trend toward a more severe behavioral phenotype **(Fig. D** to **H)**. A principal component analysis (PCA) of all ASD-like symptoms showed that Treg stimulation or depletion consistently prevented or exacerbated, respectively, the ASD phenotypes: the first PCA axis (which captures most of the variance) revealed that the MIA_F1_ group was far apart from that of Ctrl_F1_, while the DTR-MIA_F1_ one was even further apart **(Fig 1I)**.

To better address the ecological and ethological behavioral aspects of the mice, we next used the live mouse tracker (LMT) device, which automatically tracks, identifies, and characterizes the social interactions of mice over 14 h under more physiological conditions (*22*). Using this, we generated a comprehensive behavioral profile of each mouse, comparing the MIA_F1_ to the Ctrl_F1_ mice **(Fig. 1, J** to **N)**. We found that adult PBS-MIA_F1_ mice had fewer contacts of all types (side-to-side, opposite side-to-side, oral–oral, oral–genital) **(Fig. 1, J** and **K)**. In contrast, IL2-MIA_F1_ showed improvements in all these social behavioral deficits, reaching significance for time moving in contact (**Fig 1J)** and oral–oral contact **(Fig 1K)**. In terms of social configuration, PBS-MIA_F1_ adult mice showed reduced formation of social groups of two and three, and (to a lesser degree) of behaviors and social approach **(Fig. 1, L** and **M)**. In turn, these abnormalities were rescued in IL2-MIA_F1_ mice; indeed, IL2-MIA_F1_ mice even formed more triads and had more social approaches than the PBS-Ctrl_F1_ **(Fig. 1, L** and **M).** Finally, whereas PBS-MIA_F1_ mice moved more frequently in isolation, IL2-MIA_F1_ mice were partially rescued for this phenotype, and they spent less time immobile in isolation **(Fig. 1N)**. Of note, maternal Treg depletion or stimulation alone did not induce ASD phenotypes in the offspring **(Fig. S1)**.

Taken together, we showed that maternal Tregs are critical modulators for triggering _MIA_ASD. As Treg stimulation led to blockade of IL-17A production in MIA-induced dams, these results are consistent with a triggering role of IL-17A in _MIA_ASD and raise the interesting possibility that ASD in humans could be prevented by stimulating Tregs in pregnant mothers with MIA.

### MIA imprints a long-lasting quantitative and functional defect in offspring Tregs

Similar to our observation that individuals with ASD have reduced numbers of Tregs (*15*), we observed a decrease in the percentage of Tregs in adult MIA_F1_ mice **(Fig. 2A).** As MIA can lead to epigenetic modifications of T cells in the fetus (*23*, *24*), we next analyzed the methylome of Tregs **(Fig. 2B** to **D**). We found that Tregs from PBS-MIA_F1_ had 52 differentially methylated regions (DMR) as compared to PBS-Ctrl_F1_ **(Fig. 2B).** The heatmap of significantly modulated DMRs could be separated into two distinct modules that efficiently distinguished MIA_F1_ from Ctrl_F1_ (**Fig. 2C**). Regions with reduced DNA methylation in MIA_F1_ Tregs converged towards brain development pathways, whereas regions enriched for DNA methylation converged towards lymphocyte differentiation and function pathways (**Fig. S2**). There was no significant variation in the Treg-specific demethylated region (TDSR), a non-coding region of the *FoxP3* gene locus (*25*). PCA confirmed the good classification performance of the DMR, with a perfect separation of the two conditions (**Fig. 2D**), further supporting the relevance of these methylome changes.

**Fig. 2.**
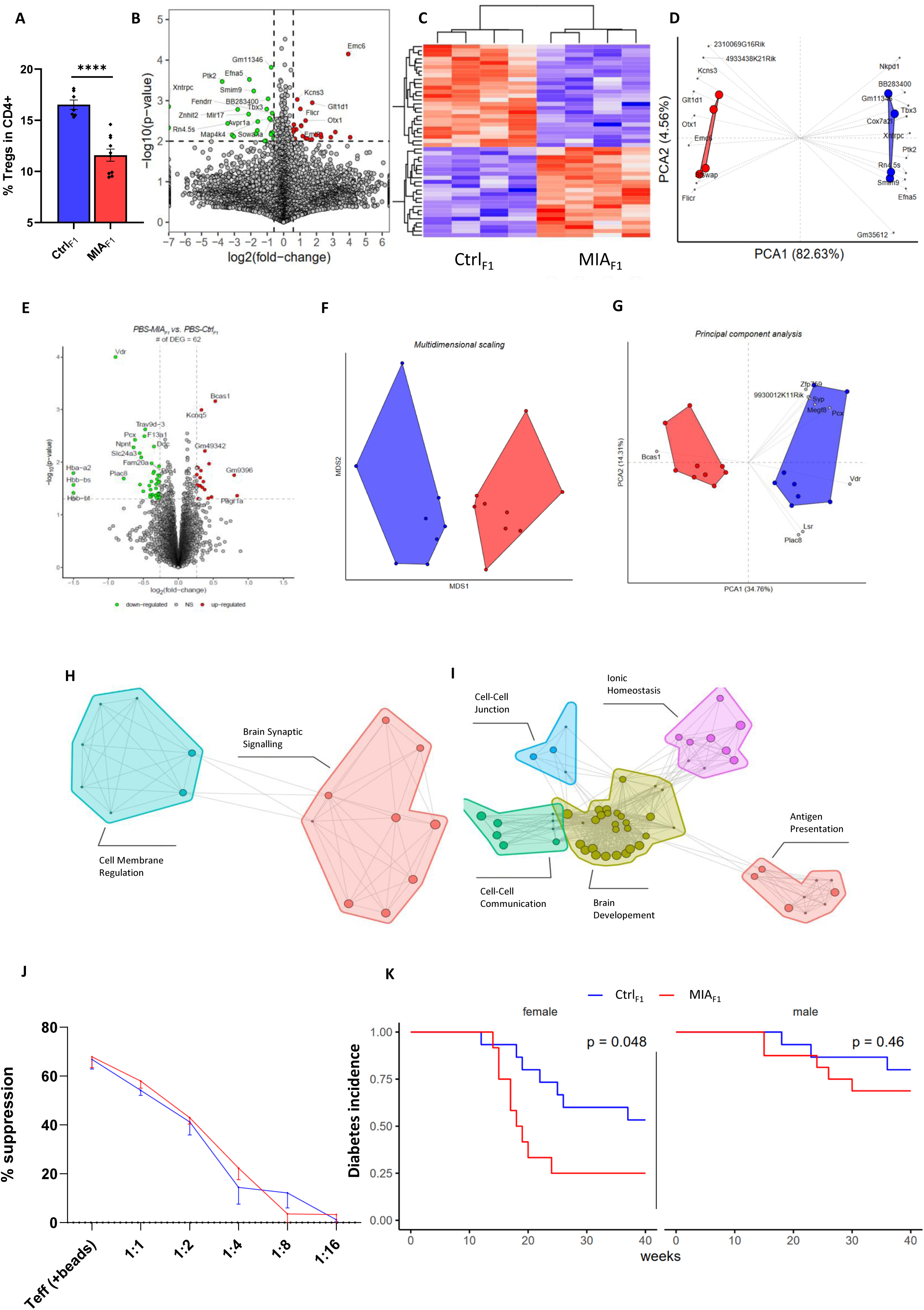
MIA induces long-term quantitative and functional abnormalities in MIA_F1_Tregs. (**A**) Flow cytometry analysis of spleen Tregs from PBS-Ctrl_F1_ (*n* = 7, 3 experiments) or PBS-MIA_F1_ (*n* = 10, 4 experiments). **(B** and **C)** Methylomic analysis of spleen Tregs from PBS-Ctrl_F1_ or PBS-MIA_F1_, represented as volcano plot of the differentially methylated region (DMR) (B) and heatmap of the DMR (C). Red, up-methylated region; blue, down-methylated region, as compared to PBS-Ctrl_F1_. **(D)** PCA of the DMR. Representation of genes shows a distance related to an origin higher than 0.85. **(E** to **G)** Bulk transcriptomic of spleen Tregs from PBS-Ctrl_F1_ and PBS-MIA_F1_, represented as volcano plot of downregulated genes (yellow) and upregulated genes (red) (E), with multidimensional scaling (MDS) classification (F), and PCA and correlation circle (G). **(H** and **I)** Comparison of differentially expressed genes (DEGs) in PBS-MIA_F1_ as compared to PBS-Ctrl_F1_ with respect to gene ontology of the cellular functions (H) or biological functions (I). **(K)** In vitro suppressive assay with spleen Tregs from PBS-Ctrl_F1_ and PBS-MIA_F1_ (*n* = 3 mice per group, with 3 replicates per group). **(L)** Incidence of diabetes in PBS-Ctrl_F1_ and PBS-MIA_F1_ from NOD mice. Left panel: females; right panel: males. Statistical analysis, t-test; \*\**P* < 0.05, \*\**P* < 0.01, \*\*\**P* < 0.001, \*\*\*\**P* < 0.0001. Results are presented as means +/– SEM.

To gain further insight into the effects of MIA on offspring Tregs, we next performed bulk RNA sequencing (RNA-seq) on sorted spleen Tregs (**Fig. 2, E** to **G**). We found only 62 differentially expressed genes (DEGs) in Tregs from PBS-MIA_F1_ as compared to PBS-Ctrl_F1_ (**Fig. 2E** and fig. S3). The relevance of these DEGs was confirmed by multidimensional scaling (MDS) (**Fig. 2F**) and PCA (**Fig. 2G**), both of which accurately classified the Tregs transcriptome as being from MIA_F1_ or Ctrl_F1_. Of note, the separation on the first PCA axis for MIA_F1_ Tregs was largely driven by the DEGs *Vdr* (encoding the vitamin D receptor), which was downregulated, and *Bcas1* (encoding the brain-enriched myelin associated protein 1), which was upregulated (**Fig. 2G**). Vdr is a key regulator of Treg differentiation, and its polymorphism has been associated with ASD (*26*); thus, low *Vdr* expression may be a mechanistic driver of MIA_F1_ Treg deficiency, as previously shown for arthritis (*27*). Consistently, vitamin D deficiency associates with the development of autoimmune diseases, such as multiple sclerosis and type 1 diabetes, and vitamin D supplementation reduced the incidence of autoimmune diseases in a large clinical study (*28*). Further studies should investigate *Vdr* as a potential key gene associated with Treg insufficiency in MIA_F1_ and its therapeutic implications.

*Bcas1* is expressed in immature oligodendrocytes (*29*) and is also a candidate gene in ASD (*30*). Of note, we found a trend for increased Olig2 staining in the brain of MIA_F1_, suggesting that MIA_F1_ oligodendrocytes have an immature phenotype (**Fig. S4).** Since Tregs promote oligodendrocyte differentiation (*13*), and tissue Tregs express genes related to their environment, the increased *Bcas1* expression could be a marker of a chronic brain inflammation. Along the same line, gene ontology (GO) analysis of the global changes in the MIA_F1_ Treg transcriptome revealed that, at the cellular component level, MIA impaired pathways related to synaptic signaling in the brain (**Fig. 2H).** At the biological pathway level, MIA disrupted several neurodevelopmental pathways **(Fig. 2I)**.

In a classical in vitro suppression assay, we did not observe a marked functional deficiency of Tregs from MIA_F1_ as compared to Ctrl_F1_ (**Fig. 2J**). However, this assay mainly captures the capacity of Tregs to deprive effector T cells of IL-2. Indeed, a more comprehensive in vivo assessment of their suppressive function revealed a clear defect in Tregs. Specifically, we performed a MIA in pregnant, non-obese diabetic (NOD) mice, whose susceptibility to spontaneous diabetes has been linked to a Treg defect (*31*). Female (but not male) NOD MIA_F1_ developed diabetes significantly faster and more frequently than NOD Ctrl_F1_ (note that NOD males are more resistant to diabetes than NOD females) (**Fig. 2K**). This experiment clearly demonstrated that MIA imprints a Treg insufficiency in the offspring. Overall, MIA induces an epigenetically imprinted global functional impairment of Tregs.

### MIA induces a meningeal pro-inflammatory state in offspring that is mitigated by Treg stimulation

Meningeal lymphocyte populations are now recognized as important players in brain development, homeostasis, and cognitive function (*32*). Of the few (still poorly-defined) resident cells in the meninges, IL-17–producing cells appear to be of particular importance for proper brain development and function (*33–35*). We thus analyzed the meningeal immune microenvironment in MIA_F1_ adult mice, treated or not with IL-2. For this, we analyzed harvested meningeal cells by single-cell RNA sequencing (scRNA-seq) and flow cytometry.

scRNA-seq of CD45^+^ cells **(fig. S5)** revealed that MIA significantly increased the meningeal monocyte and neutrophil populations (**Fig. 3A)** and altered their transcriptional programs **(Fig. 3, B** and **C).** Of note, monocyte and neutrophil recruitment to sites of inflammation is known to be IL-17 driven (*36*). The increased numbers of monocytes persisted in IL2-MIA_F1_ (**Fig. 3A**), while their transcriptome was partially restored (**Fig. 3B)**. In contrast, in IL2-MIA_F1_, the neutrophil numbers returned to normal in IL2-MIA_F1_ (**Fig. 3A**), and their transcriptomes were fully restored to the expression levels of Ctrl_F1_ (**Fig. 3C**). Functional enrichment analysis based on the GO database highlighted that MIA_F1_ have an inflammatory meningeal microenvironment, with increased pro-inflammatory signaling, reactive oxygen species production, and activation of pro-apoptotic signaling (**fig. S6A)**, associated with decreased antigen presentation and immune cell migration pathways **(fig. S6B).** Critically, IL-2_LD_ treatment reverted this inflammatory phenotype to a normal state (**Fig. S6, C** and **D).**

**Fig. 3.**
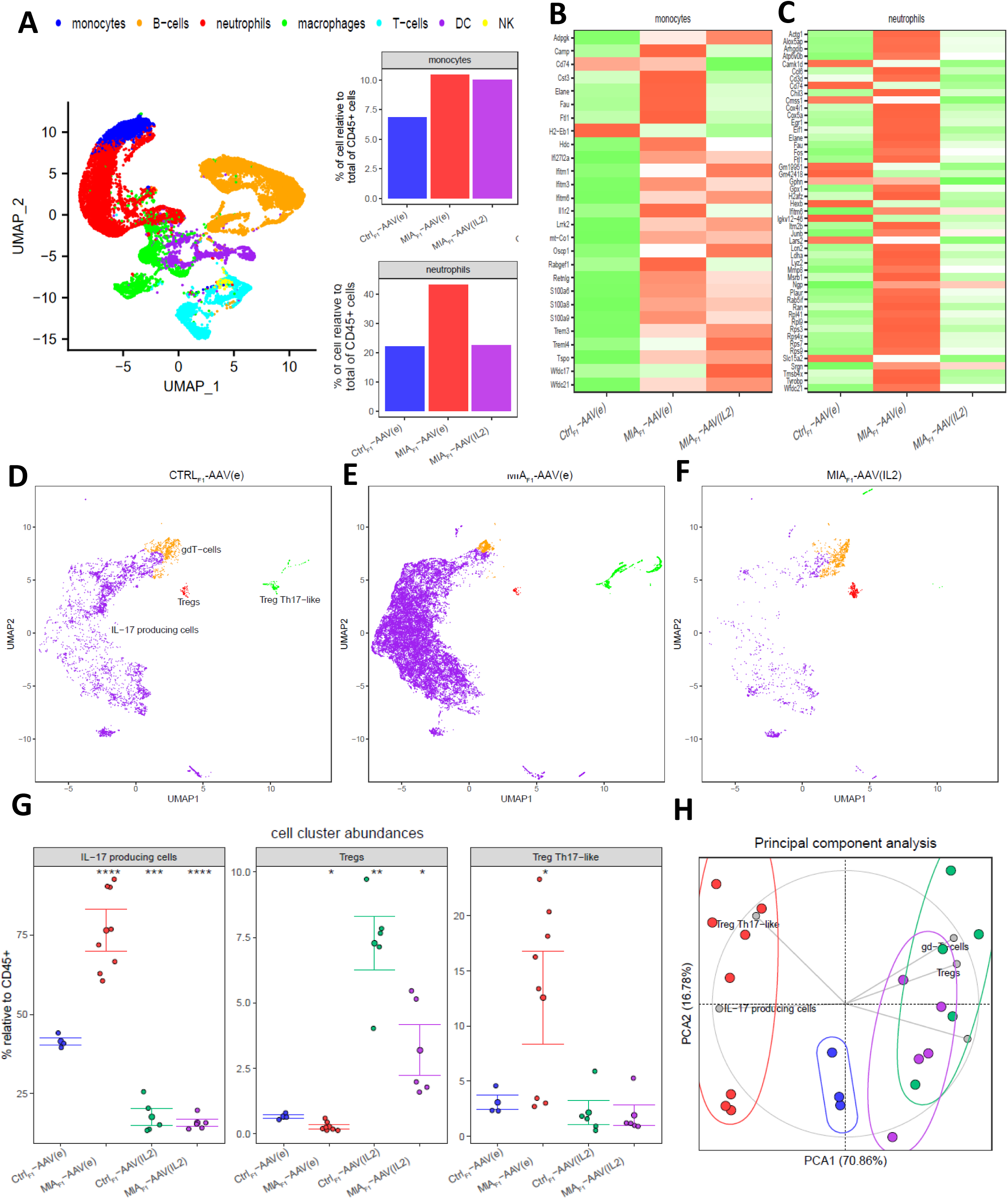
MIA induces a meningeal inflammatory microenvironment that is rescued by Treg stimulation. **(A)** UMAP representation of the single-cell transcriptomics from CD45+ meningeal cells (left panel). Percentage of monocytes and neutrophils relative to total CD45+ are shown for the different conditions. **(B** and **C)** Heatmaps of monocytes (B) or neutrophils (C) of differentially expressed genes. **(D** to **F)** UMAPs of the unsupervised flow cytometry analysis of meningeal cells from adult mice of Ctrl_F1_ (D), MIA_F1_ (E), or MIA_F1_ with long-term IL-2 exposure (F); cell cluster abundances (G) and the PCA representation of the different conditions (H) are shown. \**P* < 0.05, \*\**P* < 0.01, \*\*\**P* < 0.001, \*\*\*\**P* < 0.0001. Results are presented as means +/– SEM.

To gain more insight into the phenotype of the MIA_F1_ T cells, we analyzed CD45+ meningeal cells by flow cytometry, using unsupervised analyses to limit biases in cell population gating **(Fig. 3D to G and Fig. S7)**. MIA dramatically increased Rorγt+ IL-17A producing cells, as well as a small population of Treg/Th17–like cells, and decreased the small population of meningeal Tregs (**Fig. 3E and G**). IL-2_LD_ strikingly reversed this phenotype: in IL-2_LD_–treated MIA_F1_ mice, IL-17–producing cells and Treg/Th17-like cells populations were almost completely wiped out, and Tregs were increased **(Fig. 3F and G)**. Finally, a PCA analyses of the flow cytometry results confirmed that Tregs and IL-17–producing cells are central to accurately distinguishing MIA_F1_ from Ctrl_F1_ **(Fig. 3H).**

Meninges were recently found to host Tregs (termed mTregs) (*37*). Here, we found that mTregs were activated and expanded by a systemic treatment with IL-2_LD_ of Ctrl _F1_ and MIA_F1_ mice **(fig. S8A** to **C).** More specifically, IL-2_LD_ expanded the mTreg clusters 3 and 8 in the single-cell analyses focused on lymphocytes **(fig. S8B).** This population is characterized by a thymic origin (Helios^+^), expression of canonical Tregs markers (CD4^+^, CD25^+^, Foxp3^+^), a resident signature (CD69^+^, KLRG1^+^, IL1Rl1^+^) with a specific meningeal chemokine migration profile (CCR2^+^, CXCR6 ^+^) (*34*), and an activated phenotype (ICOS^+^, PD1^+^, CTLA4^+^, LAG3). This phenotype differentiate the expanded mTregs from previously characterized brain Tregs, which are CCR5^+^, CCR6^+^, and CCR8^+^ (*38*) **(fig. S8D).** mTreg from MIA_F1_ also exhibited a resident phenotype, but with an increased expression of CCR8, indicating a pro-inflammatory environment **(fig. S8E)** (*39*). Strikingly, the mTregs from IL-2_LD_–treated MIA_F1_ displayed an increased expression of *Foxp3*, indicative of a more suppressive phenotype **(fig. S8 F**) and of *Spp1* (osteopontin) **(fig. S8F**), which is expressed in a brain Treg population that controls microglia reactivity and promotes oligodendrocyte maturation (*40*). Finally, a T cell antigen receptor (TCR) repertoire study of the single-cell transcriptomes showed a polyclonal repertoire with no expansions **(fig. S8G).** In summary, meningeal cells of F1 mice with MIA-related ASD displayed a chronic pro-inflammatory environment, associated with a major increase of IL-17A–producing cells and neutrophils, and a decrease of Tregs—all of which were rescued by IL-2_LD_ treatment.

This meningeal inflammatory state was accompanied by changes in the brain tissues, as evidenced by histology and imaging. Histological studies showed that brain microglia staining increased in MIA_F1_ but was normalized in IL-2_LD_-treated MIA_F1_ **(Fig. 4, A and B,** and **fig. S9)**, consistent with a previous report suggesting that MIA_F1_ mice have a chronic brain inflammation (*9*). To obtain morphological insight into the MIA effects on brain, we analyzed the mice using brain 11.7-tesla magnetic resonance imaging **(Fig. 4C** and **fig. S10)**. First, we observed that MIA induced a statistically significant increase in the volume dispersion in all ten brain regions in MIA_F1_ as compared to Ctrl_F1_ mice, and mainly in the insula (agranular insular area and gustatory area), basal ganglia (striatum, amygdala, hypothalamus, claustrum), and anterior cingulate area **(fig. S10)**. Notably, these findings were normalized by IL-2_LD_ treatment of the MIA_F1_ mice. Moreover, these regions are involved in social symptoms in individuals with ASD (*41*). Second, we found that the temporal cortex volume (ectorhinal and perirhinal areas) was increased in MIA_F1_, but normalized by Treg stimulation in IL2-MIA_F1_ **(Fig. 4, E and F).** These results are in line with human brain imaging studies that found an increased temporal size in individuals with ASD (*42*). Of note, temporal volume is associated with social and communication symptoms (*43*), and both rare and common ASD-associated gene variation are expressed in the temporal lobe (*44*). Lastly, a PCA based on the volume of brain regions showed a good classification performance, driven by the volumes of the temporal cortex, anterior cingulate area, dorsal striatum, and insula **(Fig. 4D)**, in line with the literature reporting the importance of these regions in driving ASD-like symptoms (*41*, *42*). This restoration of brain structure highlights the therapeutic potential of IL-2_LD_ treatment in _MIA_ASD.

**Figure 4:**
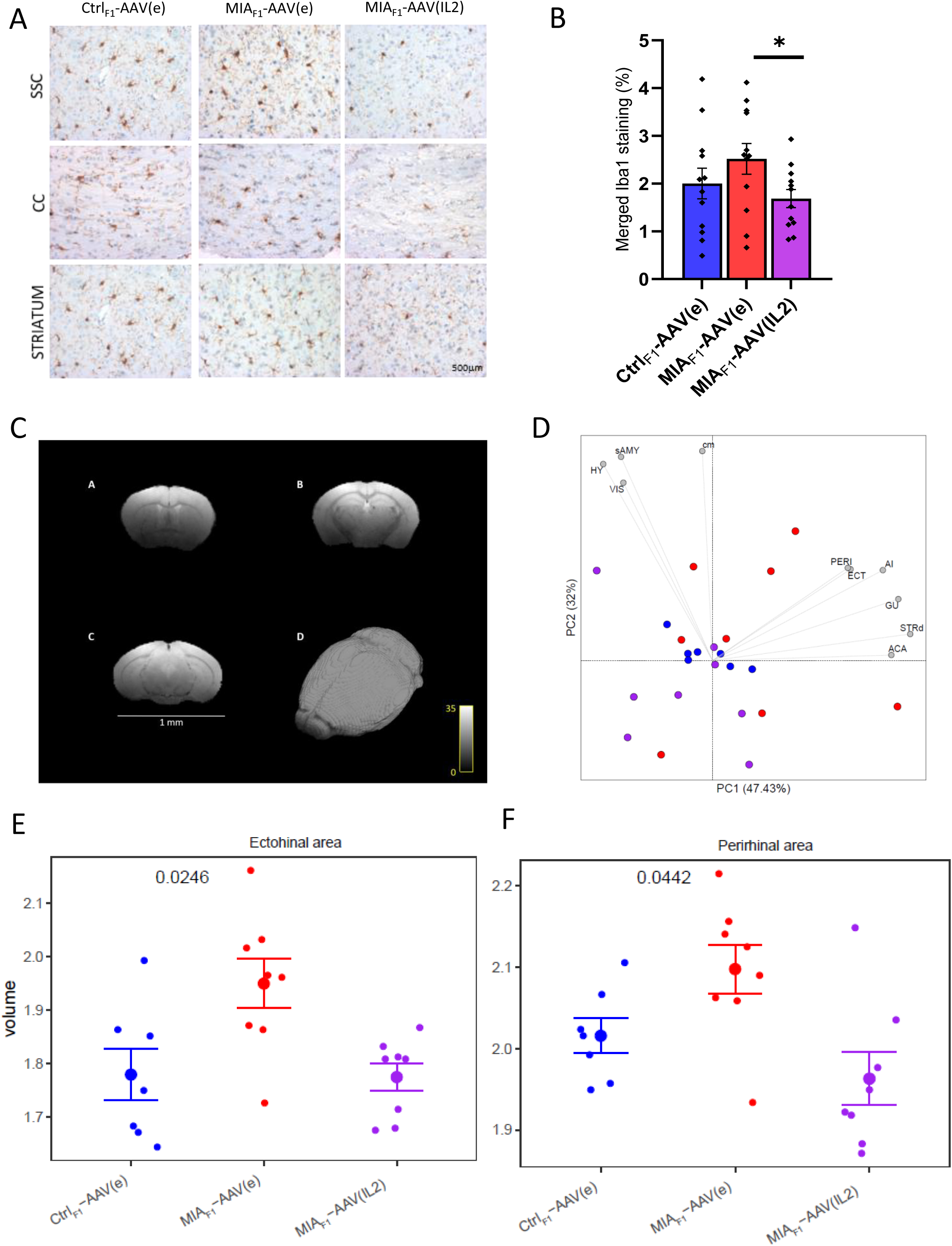
M**a**ternal **immune activation disrupts brain morphology rescued by Tregs stimulation. (A** and **B)** Brain Iba1 staining (A) in the somato sensorial cortex (SSC), the corpus callosum (CC), and the striatum, with barplot representation (B). **(C)** Brain structure from the 11.7T MRI. **(D)** PCA representations based on brain volume area. **(E** and **F)** Barplot representation of the volume. ACA, anterior cingulate area; AI, agranular insula area; CM, claustrum; ECT, ectorhinal area; GU, gustative area; HY, hypothalamus; PERI, perirhinal area; sAMY, striaum-like amygdalar nuclei; STRd, striatum dorsal area; VIS, visceral area. Statistical analysis, t-test. \**P* < 0.05, \*\**P* < 0.01, \*\*\**P* < 0.001, \*\*\*\**P* < 0.0001. Results are presented as means +/- SEM.

### Stimulation of Tregs in MIA_F1_ young adult mice cures their ASD-like phenotype

Given our results on the functional alterations of Tregs, their association with meningeal and brain inflammation, and their normalization by IL-2_LD_ treatment, we aimed to evaluate whether Treg stimulation with IL-2_LD_ would rescue the ASD-like phenotype of MIA_F1_ mice. To reduce the stress of repeated injections, we performed a single injection of an adeno-associated virus expressing IL-2 (AAV-IL2) for long-term Treg stimulation (*45*) **(Fig. 5A)**. AAV-IL2 stimulated Tregs (**Fig. 5B)**, and reduced the Th17/Treg ratio **(Fig. 5C)**, in both MIA_F1_ and Ctrl_F1_ young adult mice. Strikingly, Treg stimulation of MIA_F1_ rescued repetitive behaviors **(Fig. 5D)** and the loss of social preference **(Fig. 5E)**; these observations were also supported by PCA analysis **(Fig. 5F)**. Using the LMT device, we observed that IL-2_LD_ exposure normalized the number of contacts **(Fig. 5G)**, the social configurations **(Fig. 5H),** and the social approaches **(Fig. 5I)**, of MIA_F1_ mice. Thus, Treg stimulation corrected the altered behaviors of MIA_F1_ and mitigated brain and meningeal inflammation **(Fig. 3).** In sum, stimulation of MIA_F1_ Tregs with IL-2_LD_ cured established autistic-like behaviors, highlighting a causal relationship between Treg defects and these behaviors.

**Fig. 5.**
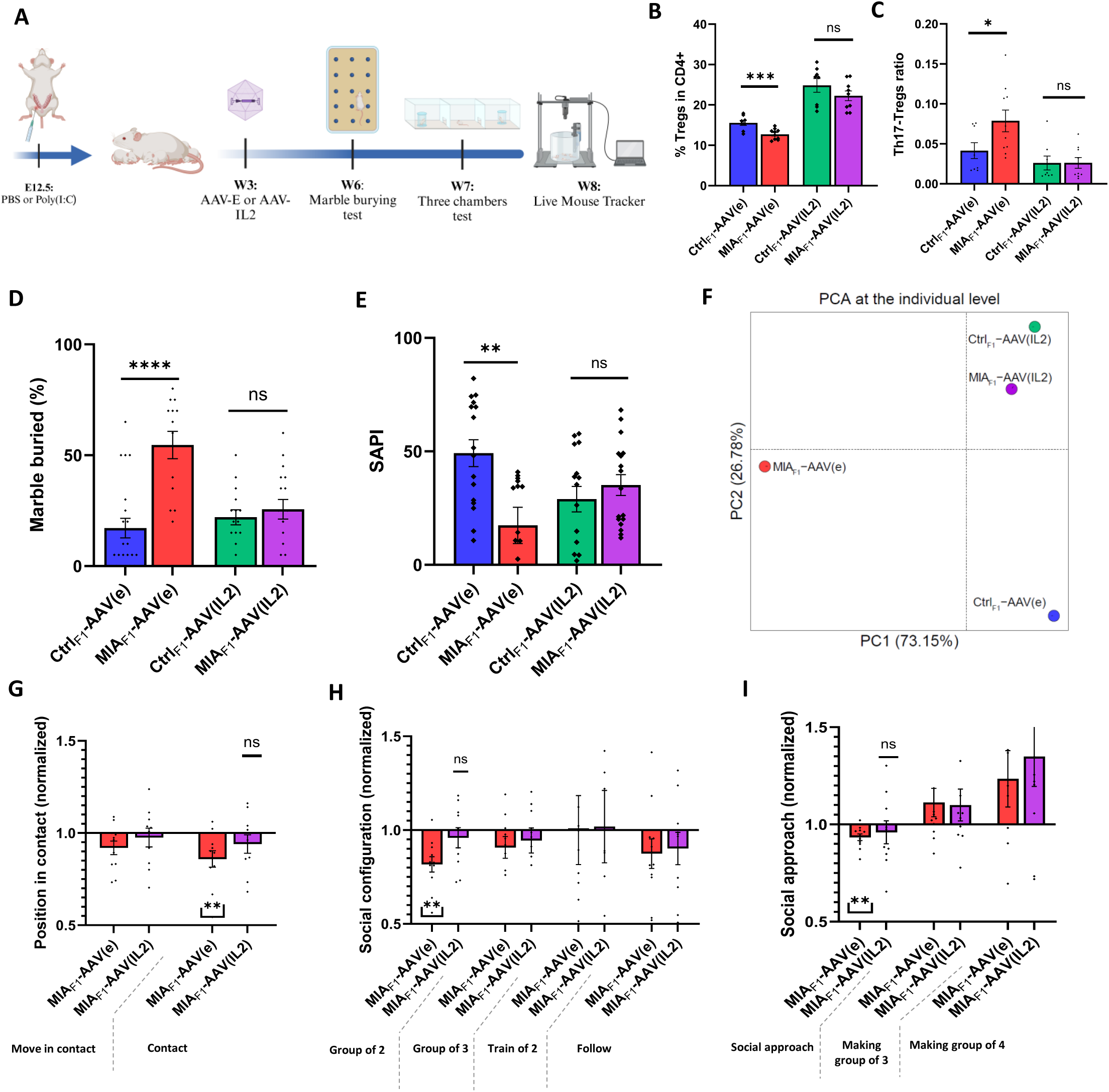
Treatment with IL2_LD_ of MIA_F1_ young-adult mice rescues autistic symptoms. **(A)** Schematic of experimental design. At week 3 (W3), pups were intraperitoneal injected with 1 × 10^10^ infectious particles of AAV(IL2) or AAV(empty) vectors. The marble burying test were evaluated at W6. After one-week isolation, mice performed the three-chamber test at W7 and were analyzed by the live mouse tracker for 14 h at W8. Some mice were sacrificed, and spleen and meningeal cells were analyzed. **(B)** Flow cytometry of Tregs at W5. **(C)** Th17/Tregs ratio at W5**. (D)** Percentage of marbles buried for the PBS_F1_-AAV(e) (*n* = 21, 4 experiments), MIA_F1_-AAV(e) (*n* = 13, 3 experiments), PBS_F1_-AAV(IL2) (*n* = 15, 4 experiments), and MIA_F1_-AAV(e) (*n* = 17, 4 experiments) **(E)** Social preference index for the following mice: PBS_F1_-AAV(e) (*n* = 21, 4 experiments), MIA_F1_-AAV(e) (*n* = 13, 3 experiments), PBS_F1_-AAV(IL2) (*n* = 15, 4 experiments), and MIA_F1_-AAV(e) (*n* = 17, 4 experiments). **(F)** PCA of the different conditions. (**G to I)** Live mouse tracker behavioral profiles for the MIA_F1_-AAV(e) (*n* = 11) or MIA_F1_-AAV(IL2) (*n* = 10), normalized to the appropriate control mice (*n* = 12 per condition) over 14 h of free interactions, showing position in contact (G), social configuration (H), and social approach (I). Statistical analysis, t-test except for LMT: Wilcoxon matched-pairs signed rank test. \*\**P* < 0.05, \*\**P* < 0.01, \*\*\**P* < 0.001, \*\*\*\**P* < 0.0001. Results are presented as means +/- SEM

## Discussion

We and others have reported that persons with ASD have an altered Th17/Treg balance, with a quantitative Treg deficit, in the peripheral blood, without evidence of a functional Treg deficiency (*15*). Here, we report an altered phenotype of MIA_F1_ Tregs, which could be explained at least in part by epigenetic imprinting, and we demonstrated a true functional Treg deficiency in vivo. Assessing the impact of this Treg deficiency on the brain microenvironment, we have previously shown that MIA leads to a highly inflammatory brain environment (*36*). We now establish a causal link between Treg deficiency and ASD, by showing that Treg stimulation can normalize both the biological and the clinical symptoms in ASD-like young adult mice. This is remarkable because it shows that a Treg defect is not only important for initiating the autistic core symptoms during prenatal growth but is also involved in the maintenance of these symptoms in the adult offspring.

Our results have important clinical applications. At the maternal level, Treg stimulation could be beneficial in limiting the increased risk of ASD following a MIA. As IL-2_LD_ is very well tolerated (*46*), the effect of a supplementation with IL-2_LD_ in case of a documented MIA should be evaluated. At the level of individuals with ASD, our results point to ASD as a chronic inflammatory brain disorder, rather than a permanently-fixed neurodevelopment state, and—importantly—one that can be reversed. This paves the way for the evaluation of Treg stimulation by IL-2_LD_ for the treatment of individuals with ASD. We propose a paradigm shift from viewing ASD as a purely neurodevelopmental disorder to viewing ASD as, at least in part, a chronic inflammatory disorder of the brain that is potentially reversible.

## Acknowledgments

We would like to acknowledge all personnel of UMS 28, and particularly Soizic Jezequel and Yohann Bertelle, for providing excellent care to the mice used in this work. We thank the RNA-seq core facility and the Preclinical Brain platform of the Paris Brain Institute, and particularly Matthieu Santin, as well as Benedicte Hoareau from the CyPS core facility.

## Funding

This work was supported by a Laboratory of Excellence grant (n° ANR-11-IDEX-0004-02) and the Janssen Horizon program. PE was supported by an INSERM “poste d’accueil” and the Robert Debré Child Brain Institute (n° ANR-11ANR-23-IAIIU-0010).

## Author contributions

Conceptualization: DK conceptualized the global project, with the participation of PE

Investigation: Investigations were conducted by PE, GF, NT, VM, NC, SB, GM, and TM

Visualization: Data visualization was performed by PE, NT, VM, TM and DK

Formal analysis: NT performed the statistical analyses

Funding acquisition: DK

Project administration: DS, RD, BY, MR and DK administered the work in their respective areas.

Validation: Results of experiments were validated by DS, RD, BY, MR, DK for the work in their respective areas.

Supervision: DK carried out the overall supervision of the work.

Writing – original draft: PE, DK

Writing – review & editing: all authors

## Competing interests

DK, MR, PE, and RD are inventors of patents hold by their institutions claiming the therapeutic use of IL-2; DK and MR hold shares in ILTOO pharma, the licensee of these patents.

## Data and materials availability

All data, code, and materials used in the analysis are available upon request.

## List of Supplementary Materials

## Materials and Methods

figs. S1 to S10

