## Supplementary for "Maternal immune activation imprints a regulatory T cell deficiency in offspring that drives an autism-like phenotype"

### Materials and methods

Animals

All experiments were performed according to the guidelines established by the European Community Council (Directive 2010/63/EU) and approved by the ethics committee (APAFIS #17990). Animals were maintained on a 12h light-dark schedule in a temperature (22+/-1°C) and humidity-controlled (50%) environment. Food and water were provided ad libitum. C57BL/6-Foxp3-EGFP transgenic mice expressing GFP under the control of the Foxp3 gene promoter were provided by B. Malissen (Luminy, Marseille, France). Foxp3-DTR (human diphtheria toxin receptor) C57BL/6 mice lead to specific depletion of Tregs by using diphtheria toxin (DT) injection due to expression of DT receptor under the control of the Foxp3 promoter (*1*) were purchased from Jackson lab. Non-obese diabetic (NOD) mice have a susceptibility to spontaneous diabetes linked to a defect in Tregs (*2*) were provided by V.Kuchroo (Brigham and Women's Hospital, Boston, MA).

Maternal immune activation

Mice were mated overnight and females were checked the day after for the presence of seminal plugs, (Day 0.5 – E0.5). On E10.5, female mice were weighed. On E12.5, pregnant mice were injected with a single dose (5mg/kg, i.p) of Poly(I:C) (Sigma Aldrich - P9582) or the same volume of PBS. Each dam was returned to its cage and left undisturbed until birth. All pups remained with their mother until weaning on W3 at which time males were group-housed at a maximum of 5 per cage with a littermate.

Maternal Tregs stimulation

Between gestational day 7.5 to 11.5 pregnant mothers were subcutaneously injected with either PBS (used as a control of injection) or subcutaneous injections with 50,000 units of ILT-101 (human recombinant IL-2; ILTOO Pharma) during 5 consecutive days and compared with an injection of PBS.

Maternal Tregs depletion

To deplete Tregs in mothers, we used Foxp3-DTR mice. In these mice Diphtheria toxin (DT) injection mediated specific elimination of Tregs. We injected i.p pregnant mice twice with DT (50µg/kg) on gestation days 10.5 and 11.5.

ELISA

Three hours post-Poly (I:C) injection, serum preparation, and IL-17A protein level measurements were performed according to manufacturer's manual (Invitrogen).

Monitoring Type 1 diabetes incidence

We applied the maternal immune activation protocol described previously to NOD females. Offspring from PBS or Poly(I:C) injected mice were then monitored once a week for diabetes incidence using urine dipsticks.

Offspring Tregs stimulation

At W3 post-natal, IL-$2_{LD}$ was administered by a single injection of an IL-2–producing recombinant AAV vector serotype 8 (AAV8) (rAAV) vector and compared with the injection empty rAAV, both at a concentration of $10^{10}$ rAAV viral genomes as previously described (24). AAV verification was performed by blood sampling and Treg analysis at W5.

Behavioral analysis

Before all experiments, the mice were transported to the behavioral room and given one hour to habituate. Only males were tested (except for ultrasonic vocalizations) and all behavioral tests (except for the Live Mouse Tracker) were carried out during daylight hours, between 9 am and 6 pm, under dim light, in a quiet environment. Behavioral tests were conducted in the following order: marble burying test (at week 6), three-chamber (at week 7), followed by live mouse tracker (between week 8 and week 10). Between two behavioral tests, the mice were left for at least 5 days.

Ultrasonic vocalization

USV were recorded in both male and female mice at P7 (*3*). The testing pup was separated from the littermates and placed in an empty container with a foam floor. The container was placed inside a Faraday box to attenuate the surrounding noise. Spontaneous ultrasonic vocalizations were recorded for five min using Avisoft UltraSoundGate Recorder system (Avisoft Bioacoustics, Glienicke, Germany; 300 kHz sampling rate, 16-bit format). After the recording, pups were weighed and immediately returned to their home cage. The vocalizations are then automatically analyzed using the specific "USV processing" module on ICY (*4*).

Marble burying test

This test was conducted to assess repetitive behavior at W6 (*5*). Twenty glass beads were placed in four rows of five beads equidistant from each other in a 25*30*40 cm cages filled with 5 cm of clean bedding. After one hour of habituation in a dimly lit room, the mice were placed in the cage for 20 minutes. At the end of the period, the mice were carefully removed and returned to their cages and the number of buried balls was recorded. Marbles buried more than 50% of the ground were considered buried.

Three-chamber test

The three-chamber test was used to assess sociability and was conducted as described previously (*6*) at W7. The cage consists of 3 equal-sized chambers with transparent walls and 1 cup in each side chamber. The experiments were evaluated in a dimly lit room. The mice were isolated one week before the experiment to increase their social appetite. In the first trial (habituation), the mice were placed in the central chamber and allowed to freely explore the 3 chambers for 10 minutes. At the end of the habituation period, the mouse was gently returned to the central chamber. A stranger mouse (of the same strain and age) and an object were then placed in each side chamber inside the cup. The stranger mouse was habituated to stay under the pot for 2 sessions of 15 minutes the day before the experiment. The placement of the object and the stranger mouse were alternated. The tested mouse was then gently replaced in the cage and allowed to freely explore for 10 minutes. A video tracking system was used (SMART) to automatically quantify the time sniffing around each cup and a social preference index was calculated as previously described (*7*): (time sniffing stranger –time sniffing empty)/(time sniffing stranger + time sniffing empty). The cage and cups were cleaned between each session.

Live Mouse tracker

We placed the experimental cage (50x50x30 cm) under the setup (70 lux, T=22°C). New fresh bedding (2-3 cm high) covered the bottom of the cage; bedding is renewed for each animal. Six groups of four mice were constituted. For each experiment, we grouped two wild-type and two experimental mice and food and water ad libitum. Recording started after 1h of habituation for 14 h between 5 pm to 8 am. We used the software Python 3.6 (Python Software Foundation. Python Language Reference, version 3.6; available at http://www.python.org) as detaied in the original publication (*8*).

Flow cytometry staining

Blood and spleen were subjected to flow cytometry using Cytoflex LX (Beckman Coulter) and the following antibodies: CD3 PE610 (BD - Clone 145-2C11) 1:400; CD4 V500 (BD – Clone RM4-5) 1:400; CD25 PeCy7 (BD - Clone PC61) 1:200. For intracellular staining, cells were permeabilized using Foxp3 kit (eBioscience) according to manufacturer manual and the following antibodies Foxp3 FITC (eBioscience – Clone Fjk-16s) 1:200 and RORgt PE (eBioscience – Clone B2D) 1:200. Data were analyzed with FlowJo v10.10.

Meningeal extraction and digestion

Mice were anesthetized with a solution of Ketamine 1000 (100mg/kg) and Xylazine (Rompun 2% 10mg/kg) (80/20) diluted in NaCl. Mice were then intracardially perfused with 50mL PBS to exsanguinate. The head was removed by cutting above the shoulders. The skull was clean from skin and flesh. Next, a cranial flap was made by cutting along the outer canthi of the eyes and between the 2 eyes. The brain was removed and the cranial flap was transferred to a petri dish with cold PBS under a binocular loupe. The meninges were removed from the inner part of the cranial flap and transferred in Ice cold buffer (1X PBS, 1% BSA, 1mM EDTA). The meninges were then digested in 2mL of 1mg/mL Collagenase VIII (sigma), 0.5mg/mL DNAse I (Sigma) in complete media (DMEM + 2 %FBS) for 15 minutes at 37°C with shaking at 130RPM. The digestion was inactivated with 10mL of Ice-cold buffer. The digested meninges were filtrated on a 70µm cell strainer and centrifugated at 400g for 4 minutes à 4°C. The cells were then stained as described above.

Unsupervised analysis of flow cytometry data

Cytometry profiles were displayed using UMAP (Uniform Manifold Approximation and Projection) representations based on the expression of all markers (*9*). This technique facilitated the reduction of data dimensionality while preserving the overall structure of cell populations, thereby enabling the visual identification of subpopulations. The K-means algorithm was used to automatically identify 150 clusters of similar cells. The quality controls of these unsupervised analyses are detailed in Supplementary Figure 7.

Cytometry profiles were visualized using UMAP (Uniform Manifold Approximation and Projection), leveraging the expression data of all markers (*9*). This approach effectively reduced the dimensionality of the dataset while retaining the global structure of cell populations, enabling clear visualization of subpopulations. To identify clusters of similar cells, the K-means algorithm was applied, resulting in 150 distinct clusters. Detailed quality control metrics and validation of these unsupervised analyses are provided in Supplementary Figure 7.

Immunohistochemistry

Mice (PBS-AVV(e) (n=4), PBS-AVV(IL2) (n=4), Poly(I:C)-AVV(e) (n=4), Poly(I:C)-AVV(IL2) (n=4)) were perfused directly intracardially with PFA. Brains were carefully removed and placed for 24h in a solution of 4% PFA. Brain tissue was embedded in paraffin and 5mm consecutive sections were performed throughout the brain. One of every ten sections was stained with hematoxylin-eosin (H-E). Immunohistochemistry with anti-GFAP (clone 6F2, mouse monoclonal, Dako, 1/500), anti-Iba1 (clone 20A12.1 mouse monoclonal, MABN92, 1/500), and anti-Olig2 (mouse monoclonal, EPIT-MICS, 1/1000) was performed using the Bench-mark station automated system (Ventana Medical system Inc., Roche). All antibodies were pretreated using a proprietary buffer pH8 (CC1) and incubated for 36 minutes (GFAP) or 64 minutes (Olig2) at 95ºC or 64 minutes at 100ºC (Iba1). For tissue analyses, regions of interest were drawn in three sections of the hippocampus located between Bregma –1.46 and –2.46mm and three sections of striatum, corpus callosum and cortex situated between Bregma 1.10 and 0.14mm. Quantification was done using the open source software for bioimage analyses: QuPath (*10*).

Statistics and reproducibility

Statistical analyses were performed using GraphPad Prism. No statistical methods were used to predetermine the sample size. The sample size was chosen based on recommendations and previously conducted studies (*6*). The normal distribution of experimental data was assumed and not formally tested. For two-group comparisons, an unpaired student's t-test was used. Differences between more than two groups were assessed using one-way ANOVA with Tukey's post hoc test. Details of statistical comparisons are listed in the relevant figure legends. All data are represented as mean +/- SEM. Each data point denotes individual animals. Significant differences are indicated in the figures by * p-value<0.05, ** p-value<0.01, *** p-value<0.001, **** p-value<0.0001.

Bulk transcriptomic

Spleen cell sorting

In Foxp3-GFP mice, spleens were harvested, diluted, and filtered on 70µM cell strainer. Cells were then magnetically enriched with biotin antibodies to Ter119, B220, CD11c, CD8a and anti-biotin micobeads (Miltenyi) (1:500). The biotin-labeled cells were then passed over a Myltenyi LS magnetic column and the negative fraction recovered. This enriched fraction was stained with a CD45 PE-TR (BD- Clone 30-F11) (1/1000) and Live/Dead EF-780 (1/1000) and CD4 V500 (1/400). Live CD45+ CD4+ Foxp3+ cells were sorted using ARIA-II (BD).

Extraction of Treg RNA and DNA

DNA and total RNA were extracted simultaneously using the AllPrep DNA/RNA/Protein kit (QIAGEN) according to the manufacturer's protocol. RNA was then processed at the Paris Brain Institute Platform.

Bulk RNAseq analysis

RNA was preserved in a frozen lysis buffer to maintain sample integrity. Extraction quality was assessed by measuring the RNA Integrity Number (RIN), with a quality threshold set at 6. Libraries were prepared using the low-input RNA Nextera XT Kit, and RNA-seq profiling was performed on an Illumina NextSeq 500 platform (75 bp, paired-end), achieving an average depth of 15 million reads per sample. Sequenced reads were trimmed for quality and adapter content using Trim Galore (*11*) and aligned to the transcriptome reference using Salmon 3 (*12*). Differential gene expression analysis was conducted with EdgeR (*13*), and functional enrichment was performed using Enrichr (*13*) based on the BioPlanet database.

Epigenetic analysis

Genomic DNA extraction and bisulfite conversion were performed using EZ DNA Methylation-Direct Kit (Zymo). Converted genomic DNA were submitted to the Hartwell Center at SJCRH for library construction and sequencing on the Illumina NovaSeq platform with 150bp paired-end and target reads of 500 million per sample. WGBS reads were trimmed by 10bp on 5' and 3' ends as well as all Illumina adapter sequences. Trimmed reads were aligned to the mm10 genome using the BSMAP v. 2.90 software. Methylation levels were called using BSMAP. Differential methylation analysis was performed by R package DSS (version 2.34) using two-group comparisons with a p-value cutoff of 0.01. Principal component analysis was carried out on the top 3,000 CpG sites with the highest variance. For WGBS, only CpG sites with coverage higher than 5 reads in all samples were included.

Single-cell RNA sequencing of meningeal

Sorting meningeal CD45+

Meninges were extracted as described previously. Meninges were digested separately and then merged by condition (4 mice per group). Cells were stained with a CD45 PE-TR (1/1000) and Live Dead EF-780 (1/1000). Live CD45+ cells were sorted using ARIA II (BD).

DNA and RNA extraction

Single-cell TCR and RNA library construction and sequencing by 10X Genomics Chromium. After isolation of CD45+ by flow cytometry from meninges, single-cell library construction was performed using Chromium Next GEM Single Cell 5’ Library & Gel Bead kit v1.1 (PN-1000165), Chromium Next GEM Chip G Single Cell Kit (PN-1000127), Chromium Single Cell 5’ Library construction kit (16 rxns) (PN-1000020), Chromium Single Cell V(D)J Enrichment kit – Mouse T Cell (PN-1000071) and Single Index kit T set A (PN-1000213) according to the manufacturer’s protocol. Briefly, single-cell suspensions from a total of 20,000 cells with barcoded gel beads and partitioning oil were loaded to Chromium Next GEM Chip G to generate single-cell gel bead-in-emulsion. Full-length cDNA along with cell barcode identifiers were PCR-amplified to generate 5’ Gene Expression (GEX) libraries and V(D)J libraries. Libraries were sequenced on a NovaSeq 6000 (Illumina) to achieve a minimum of 23,000 pair-end reads per cell for GEX and 7000 paired-end reads per cell for V(D)J. Reads were aligned using Cell Ranger v6.1.1 to the GRCm38 mouse references.

#### Single-cell RNA-seq analysis

The expression matrices resulting from reads alignment were analyzed using the R environment. Only high-quality cells were selected for subsequent analysis. The criteria for high-quality cells included a mitochondrial gene percentage below 10% and a number of unique molecular identifiers (UMI) between 200 and 15,000. Regularized negative binomial regression, based on the 3,000 most variable genes, was used to normalize the count matrix. Dimensionality reduction was performed using principal component analysis (PCA) based on the 3,000 most variable genes. Differential gene expression analysis was conducted using the Wald t-test with p-values corrected using the false discovery rate (FDR) procedure.

#### Repertoire clonality

Single-cell TCR sequencing data were processed using the “scRepertoire” R package (*14*). Clonotypes within each condition and GEX-identified cluster were divided into count fractions based on their occurrence in the repertoire (clonotypes with an occurrence of 1 in green, and 2 in purple). The number of clonotypes belonging to each count fraction was plotted.

#### MRI

### Data acquisition

Magnetic resonance imaging (MRI) acquisitions were performed using a preclinical 11.7 T MR scanner (Biospec Bruker, BioSpin, Germany) equipped with a 1H transceiver surface cryoprobe dedicated to mouse brain imaging (Biospec Bruker, BioSpin, Germany). MRI was performed in 34 mice (APAFIS #43855) ((CtrlOff-AVV(e) (n=8), CtrlOff-AVV(IL2) (n=9), MIAOff-AVV(e) (n=8), MIAOff-AVV(IL2) (n=9)) under general anesthesia using different concentrations of isoflurane mixed with oxygen and air flows of 600 mL/min. The level of isoflurane was gradually reduced to achieve light anesthesia during resting-state functional MRI (rs-fMRI) acquisitions: 3% for induction, 2% during head positioning, 1.5% during adjustments (frequency adjustment, shimming, reference pulse), 0.8-1% for MRI acquisitions. Respiratory rate was monitored and body temperature was maintained at 37 ± 1°C using a heated water circuit integrated into the cradle. The head was placed in the supine position, stereotaxically restrained by a bite bar and ear pins. For each animal, the MRI protocol consisted of the acquisition of:

(i) $T_2$-weighted ($T_2$w-MRI) structural images using a 2D multi-slice multi-echo (MSME) sequence with the following parameters: echo time (TE) = 15 ms, repetition time (TR) = 3000 ms, flip angle (FA) = 90°, bandwidth (BW) = 50 kHz, 70 slices, slice thickness = 0.250 mm, field of view (FOV) = 200*200 $mm^2$, matrix size = 160*160, in-plane spatial resolution = 0.125*0.125 $mm^2$, 1 repetition, acquisition time = 8 min

(ii) blood oxygenation level dependent (BOLD) rs-fMRI time series using a 2D gradient echo (GE) with echo planar imaging (EPI) readout: TE = 10 ms, TR = 600 ms, FA = 90°, 2 segments, BW = 400 kHz, 24 slices, slice thickness = 0.5 mm, FOV = 225*160 $mm^2$, matrix size = 90*64, in-plane spatial resolution = 0.250*0.250 $mm^2$, 750 repetitions, time resolution = 1.2 s, acquisition time = 15 min

(iii) a second BOLD rs-fMRI time series using the same GE-EPI sequence as (ii) with a reduced number of repetitions (10 repetitions, acquisition time = 13 sec) and a reverse phase encoding to further correct for EPI distortions.

### Data analysis

Due to the motion of the mouse head during MRI acquisition under light anesthesia, 29/34 MRI datasets were used for analysis. In this study, only anatomical images (T2w-MRI) were considered. Each individual T2w-MRI was normalized in the Turone Mouse Brain Template Altas (TMBTA) space (*15*) to further analyze the morphological changes in the different conditions.

First, a study template was computed using antsMultivariateTemplateConstruction2.sh from ANTs (*16*)and skullstripped using the segment function of SPM12 (*17*) based on tissue priors of the TMBTA. The skullstripped study template was then registered to the TMBTA template using antsRegistrationSyNQuick.sh function of ANTs (*16*, *18*).

Then, each individual T2w-MRI was skullstripped and registered to the study template using the same method as previously described for the study template using SPM12 and ANTs respectively. Registration of each individual T2w-MRI on the TMBTA template and atlas resulted from the combination of the transformations computed for (i) the registration of the individual subject to the study template and (ii) the registration of the study template to the TMBTA template and atlas.

Due to the spatial resolution of our images, we chose to combine brain regions from the TMBTA atlas to build a simplified atlas that contained 10 larger brain regions such as the cortex, corpus callosum,  cerebellum, hippocampus, hypothalamus, midbrain, olfactory bulb, thalamus, striatum, white matter corpus callosum excluded.

Voxel data were then scaled to zero, and their variance was compared using Welch's t-test to determine whether the condition induced changes in intra-class variability.

Suppression assay

Cells were enriched and sorted as described previously. In summary, fresh CD4+CD25- Tconv cells were isolated from spleen and superficial lymph nodes (brachial, axillary, inguinal, cervical) B6 Foxp3-GFP mice and sorted by FACS as CD4+Foxp3GFP- cells. Tconv cells were also stained with CellTrace™ Violet for proliferation monitoring and seeded at 5 x $10^4$ cells/well with anti-CD3/CD28 Dynabeads (1 bead:1 Tconv). Tregs (CD4+Foxp3GFP+) were co-cultured at ratios of 1/1, 1/2, 1/4, 1/8, and 1/16 Tconv/Tregs, and cells were analyzed 4 days later. The percentages of suppression of Tregs were calculated with 1-(% proliferation condition/% proliferation with beads only) * 100.

Supplementary figure legends

**Figure S1: Stimulation or deletion of maternal Tregs alone does not alter offspring behavior**. (A-E) Live Mouse Tracker Behavioral Profiles. Profiles of mice from dams treated with IL-$2_{LD}$ (n=12) and Foxp3-DTR dams depleted of Tregs at E12,5 (n=12) normalized on controls (n=12 per condition) over 14 hours of free interactions. The values of each behavior of the experimental mice were divided by the average of the two control mice in each experiment. Behaviors with higher expression in the experimental mice than the mean of the control mice have a value greater than one, while behaviors with lower expression in the experimental mice than the mean of the control mice have a value less than one. (A) Position in contact, (B) Type of contact, (C) Social configuration, (D) Social approach, (E) Isolated behaviors.

**Figure S2: Maternal immune activation during pregnancy induces methylomic changes in offspring Tregs.** (A) Functional enrichment of GO cellular components for hypomethylated regions. (B) Functional enrichment of GO cellular components for hypermethylated regions. (C) Functional enrichment of GO biological processes for hypomethylated regions. (D) Functional enrichment of GO biological processes for hypermethylated regions

**Figure S3: Differentially expressed gene in spleen Tregs bulk RNA seq**. (A) Heatmap representation of differentially expressed genes (DEGs) in offspring exposed to PBS or Poly(I:C), with hierarchical clustering performed at both the gene and sample levels, comparing Control and MIA groups. (B) List of DEGs, including gene names, fold changes, p-values, and functional annotations. (C) Functional enrichment analysis of downregulated DEGs using Enrichr. (D) Functional enrichment analysis of upregulated DEGs.

**Figure S4: Maternal immune activation during pregnancy may induce changes in the morphology of offspring' oligodendrocytes.** (A) Visualization in the brain mouse atlas of the region analyzed (B) Percentage of oligodendrocyte surface on the total surface in cingular cortex (C) Percentage of oligodendrocyte surface on the total surface in the striatum (D)

Percentage of oligodendrocyte surface on the total surface in merged striatum and cingular cortex.

**Figure S5: Quality control of meningeal CD45+ single-cell RNA sequencing.** (A) Proportion of mitochondrial gene content per cell across different conditions. (B) Number of genes detected per cell within each condition. (C) Number of unique molecular identifiers (UMIs) per cell across conditions. (D) Scatter plot showing the correlation between mitochondrial gene content (%) and the number of UMIs per cell, with individual cells represented as points color-coded by condition. (E) Scatter plot illustrating the correlation between the number of genes detected per cell and the number of UMIs, with points representing individual cells and colored according to their condition."

**Figure S6: Maternal immune activation induces a persistent pro-inflammatory meningeal state that is partially reversed by low-dose Interleukin-2.** (A) Biological processes gene ontology of upregulated pathways in MIA-offspring (B) Biological processes gene ontology of downregulated pathways in $MIA_{F1}$ (C) Biological processes gene ontology of upregulated pathways in $MIA_{F1}$-AAV(IL2) offspring (D) Biological processes gene ontology of downregulated pathways in $MIA_{F1}$-AAV(IL2) offspring

**Figure S7: Quality control for the unsupervised gating analysis. (**A) Number of cells per identified cluster. (B) Heatmap showing the uniformity of marker staining across clusters. (C) UMAP representation overlaid with the expression of different markers indicating staining density. (D) Heatmap showing the expression profiles of cell markers within each metacluster, along with the dendrogram of cluster classification.

**Figure S8: The meninges host a population of Tregs that control the central inflammation induced by the MIA.** (A) UMAP of the lymphocyte populations identified with sc-RNAseq (B) effect of the MIA and the low dose IL-2 on the different clusters of lymphocyte populations (C) Gating strategy for the identification of meningeal Tregs (D) Flow cytometry of meningeal

Tregs. (E-G) Phenotypes of meningeal Tregs using Sc-RNAseq (E) $Ctrl_{F1}$ (F) $MIA_{F1}$ (G) $MIA_{F1}$-AAV(IL2) (H) Number of TCR clonotype in the lymphocyte populations isolated.

**Figure S9: Low-dose Interleukin-2 induces changes in microglia morphology. (**A) Visualization in the brain mouse atlas of the region analyzed; (B) Percentage of Iba1 staining in the Somatosensorial cortex (SSC). (C) Percentage of Iba1 staining in the Cingular Cortex (CC). (D) Percentage of Iba1 staining in the striatum.

**Figure S10: Maternal immune activation induces volume dispertion, rescue by low-dose interleukin-2.** (A) Regions-of-interest analysis pipeline. T2w-MRI were used to build a study template (arrow 1, ANTs10). T2w-MRI and the study template were skullstripped (arrows 2, SPM segment12). Individual MRIs are then registered to the study template (arrow 3, ANTs) that is registered to the turone mouse brain template (TMBTA) template (arrow 4, ANTs). Finally, transformations are inverted and applied to the modified TMBTA atlas to get the 10 brain regions in the native space of each mouse. (B) Barplot representation of the dispertion of the volume across conditions

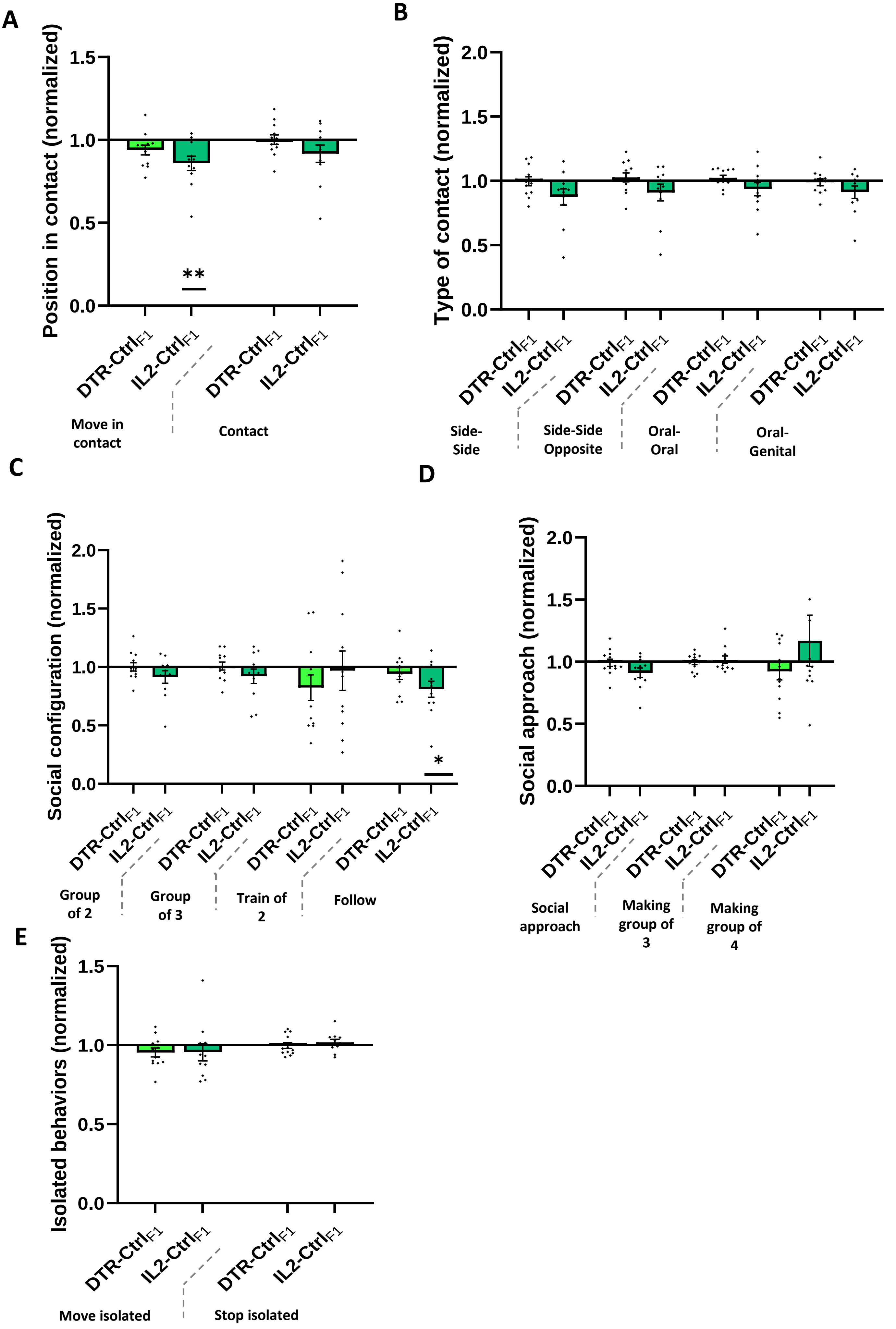

A
Position in contact (normalized)
1.5
1.0
0.5
0.0
**
DTR-Ctrl F1
IL2-Ctrl F1
DTR-Ctrl F1
IL2-Ctrl F1
Move in contact
Contact
B
Type of contact (normalized)
2.0
1.5
1.0
0.5
0.0
DTR-Ctrl F1
IL2-Ctrl F1
DTR-Ctrl F1
IL2-Ctrl F1
DTR-Ctrl F1
IL2-Ctrl F1
DTR-Ctrl F1
IL2-Ctrl F1
Side-Side
Side-Side Opposite
Oral-Oral
Oral-Genital
C
Social configuration (normalized)
2.0
1.5
1.0
0.5
0.0
*
DTR-Ctrl F1
IL2-Ctrl F1
DTR-Ctrl F1
IL2-Ctrl F1
DTR-Ctrl F1
IL2-Ctrl F1
DTR-Ctrl F1
IL2-Ctrl F1
Group of 2
Group of 3
Train of 2
Follow
D
Social approach (normalized)
2.0
1.5
1.0
0.5
0.0
DTR-Ctrl F1
IL2-Ctrl F1
DTR-Ctrl F1
IL2-Ctrl F1
DTR-Ctrl F1
IL2-Ctrl F1
Social approach
Making group of 3
Making group of 4
E
Isolated behaviors (normalized)
1.5
1.0
0.5
0.0
DTR-Ctrl F1
IL2-Ctrl F1
DTR-Ctrl F1
IL2-Ctrl F1
Move isolated
Stop isolated

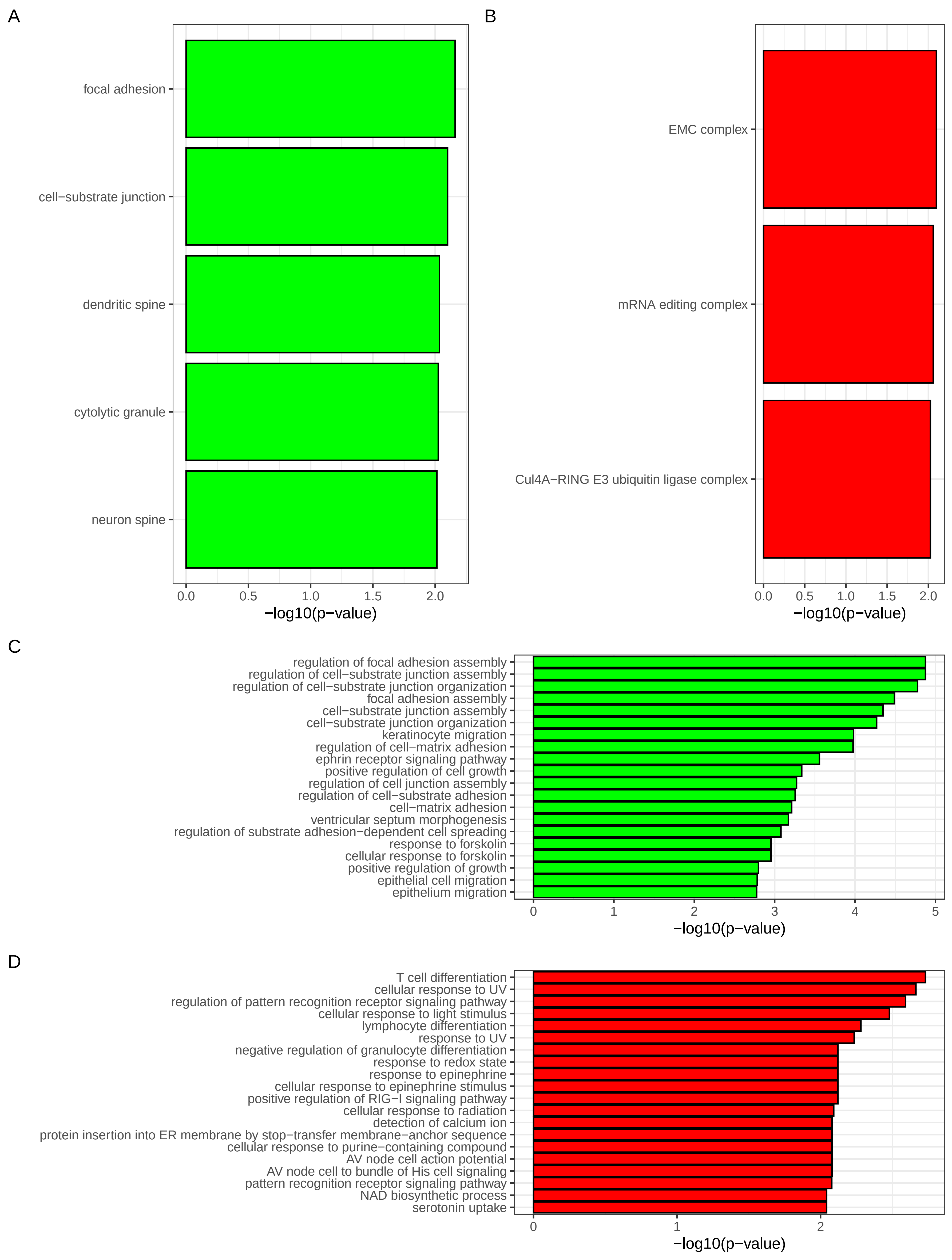
A
focal adhesion
cell−substrate junction
dendritic spine
cytolytic granule
neuron spine
0.0
0.5
1.0
1.5
2.0
−log10(p−value)
B
EMC complex
mRNA editing complex
Cul4A−RING E3 ubiquitin ligase complex
0.0
0.5
1.0
1.5
2.0
−log10(p−value)
C
regulation of focal adhesion assembly
regulation of cell−substrate junction assembly
regulation of cell−substrate junction organization
focal adhesion assembly
cell−substrate junction assembly
cell−substrate junction organization
keratinocyte migration
regulation of cell−matrix adhesion
ephrin receptor signaling pathway
positive regulation of cell growth
regulation of cell junction assembly
regulation of cell−substrate adhesion
cell−matrix adhesion
ventricular septum morphogenesis
regulation of substrate adhesion−dependent cell spreading
response to forskolin
cellular response to forskolin
positive regulation of growth
epithelial cell migration
epithelium migration
0
1
2
3
4
5
−log10(p−value)
D
T cell differentiation
cellular response to UV
regulation of pattern recognition receptor signaling pathway
cellular response to light stimulus
lymphocyte differentiation
response to UV
negative regulation of granulocyte differentiation
response to redox state
response to epinephrine
cellular response to epinephrine stimulus
positive regulation of RIG−I signaling pathway
cellular response to radiation
detection of calcium ion
protein insertion into ER membrane by stop−transfer membrane−anchor sequence
cellular response to purine−containing compound
AV node cell action potential
AV node cell to bundle of His cell signaling
pattern recognition receptor signaling pathway
NAD biosynthetic process
serotonin uptake
0
1
2
−log10(p−value)

A

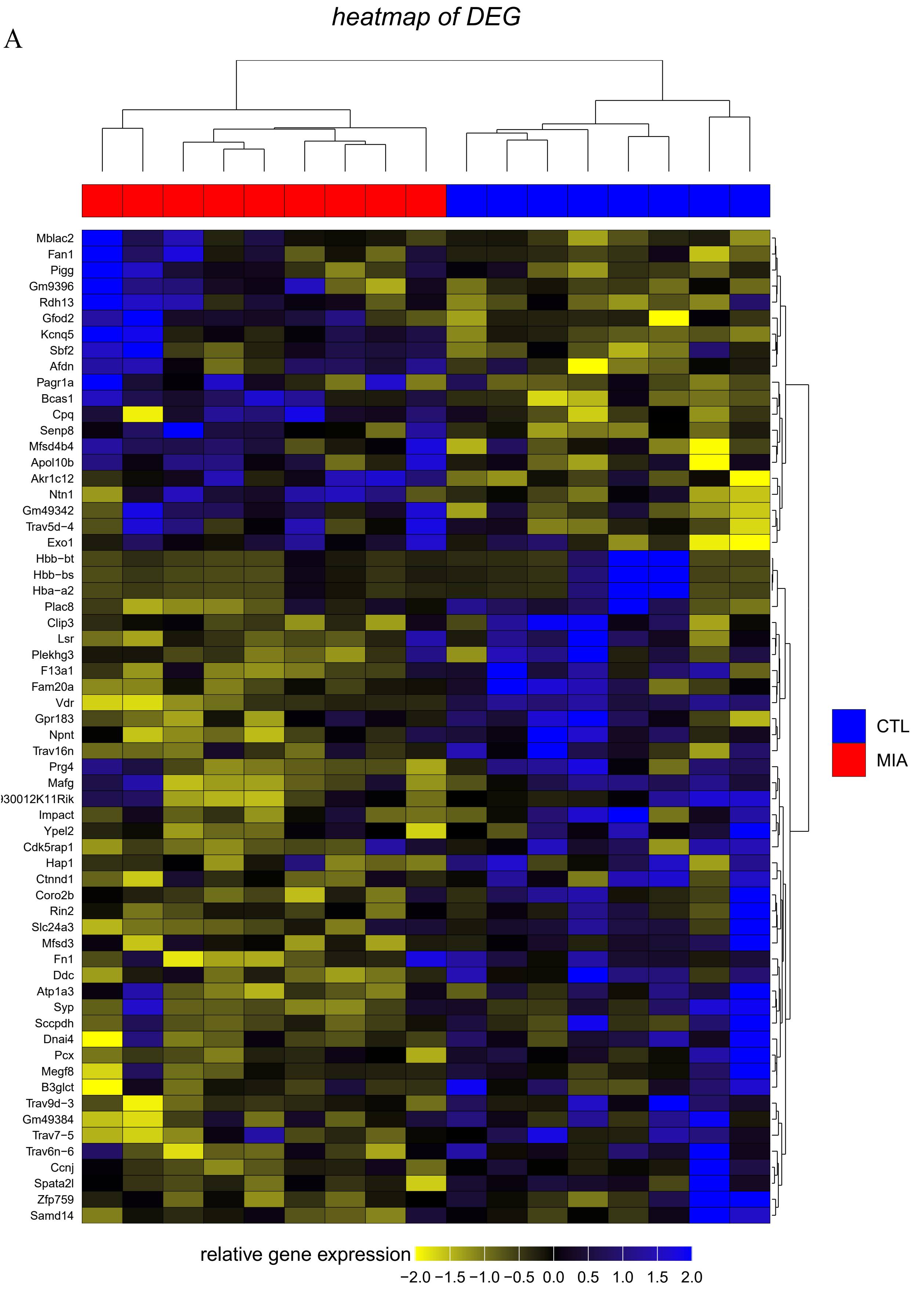


B

| GS | log2FC | p−value | description |
|---|---|---|---|
| Vdr | −0.8986773 | 9.122452e−05 | vitamin D (1,25−dihydroxyvitamin D3) receptor |
| Bcas1 | 0.5269110 | 6.915033e−04 | brain enriched myelin associated protein 1 |
| Kcnq5 | 0.3302546 | 1.010754e−03 | potassium voltage−gated channel, subfamily Q, memb |
| Trav9d−3 | −0.4750789 | 2.375265e−03 | T cell receptor alpha variable 9D−3 |
| F13a1 | −0.4852331 | 3.194118e−03 | coagulation factor XIII, A1 subunit |
| Pcx | −0.6188962 | 3.756002e−03 | pyruvate carboxylase |
| Ddc | −0.3496585 | 5.031132e−03 | dopa decarboxylase |
| Npnt | −0.6390591 | 5.232936e−03 | nephronectin |
| Gm49342 | 0.3789243 | 6.119383e−03 | predicted gene, 49342 |
| Slc24a3 | −0.5511458 | 6.731214e−03 | solute carrier family 24 (sodium/potassium/calcium |
| Fam20a | −0.5292903 | 8.059280e−03 | FAM20A, golgi associated secretory pathway pseudok |
| Prg4 | −0.3823893 | 1.047425e−02 | proteoglycan 4 (megakaryocyte stimulating factor, |
| Akr1c12 | 0.4390926 | 1.070239e−02 | aldo−keto reductase family 1, member C12 |
| Ypel2 | −0.2749340 | 1.190488e−02 | yippee like 2 |
| Mblac2 | 0.2666051 | 1.239924e−02 | metallo−beta−lactamase domain containing 2 |
| Rdh13 | 0.2819306 | 1.277644e−02 | retinol dehydrogenase 13 (all−trans and 9−cis) |
| Senp8 | 0.3225720 | 1.443939e−02 | SUMO peptidase family member, NEDD8 specific |
| Coro2b | −0.3480243 | 1.500125e−02 | coronin, actin binding protein, 2B |
| Hba−a2 | −2.0939370 | 1.611926e−02 | hemoglobin alpha, adult chain 2 |
| Trav6n−6 | −0.3573343 | 1.615748e−02 | T cell receptor alpha variable 6N−6 |
| Mfsd4b4 | 0.2637325 | 1.719825e−02 | major facilitator superfamily domain containing 4B |
| Impact | −0.2892440 | 1.758048e−02 | impact, RWD domain protein |
| Gm9396 | 0.7970663 | 1.763234e−02 | predicted gene 9396 |
| Spata2l | −0.2698101 | 1.836919e−02 | spermatogenesis associated 2−like |
| Gfod2 | 0.3107259 | 1.961229e−02 | glucose−fructose oxidoreductase domain containing |
| Plac8 | −0.7800138 | 2.044604e−02 | placenta−specific 8 |
| Ccnj | −0.3055431 | 2.092965e−02 | cyclin J |
| Trav7−5 | −0.3856538 | 2.324790e−02 | T cell receptor alpha variable 7−5 |
| Ntn1 | 0.3570920 | 2.331019e−02 | netrin 1 |
| Mafg | −0.2705103 | 2.340344e−02 | v−maf musculoaponeurotic fibrosarcoma oncogene fam |
| Dnai4 | −0.2856875 | 2.402103e−02 | dynein axonemal intermediate chain 4 |
| Ctnnd1 | −0.3165365 | 2.614763e−02 | catenin (cadherin associated protein), delta 1 |
| Cdk5rap1 | −0.5485211 | 2.652291e−02 | CDK5 regulatory subunit associated protein 1 |
| Sbf2 | 0.2677376 | 2.689339e−02 | SET binding factor 2 |
| Hbb−bs | −2.0172959 | 2.705850e−02 | hemoglobin, beta adult s chain |
| Sccpdh | −0.4145445 | 2.775966e−02 | saccharopine dehydrogenase (putative) |
| Fan1 | 0.3094619 | 2.782224e−02 | FANCD2/FANCI−associated nuclease 1 |
| Afdn | 0.2940991 | 2.813261e−02 | afadin, adherens junction formation factor |
| Cpq | 0.3419815 | 3.012549e−02 | carboxypeptidase Q |
| Clip3 | −0.3190581 | 3.130666e−02 | CAP−GLY domain containing linker protein 3 |
| Trav5d−4 | 0.3764229 | 3.305731e−02 | T cell receptor alpha variable 5D−4 |
| Lsr | −0.3965779 | 3.318224e−02 | lipolysis stimulated lipoprotein receptor |
| Samd14 | −0.5017333 | 3.563781e−02 | sterile alpha motif domain containing 14 |
| Hbb−bt | −2.1088608 | 3.830632e−02 | hemoglobin, beta adult t chain |
| Gm49384 | −0.2827424 | 3.889351e−02 | predicted gene, 49384 |
| Mfsd3 | −0.2824106 | 3.894235e−02 | major facilitator superfamily domain containing 3 |
| Trav16n | −0.2653881 | 3.922738e−02 | T cell receptor alpha variable 16n |
| Gpr183 | −0.2710990 | 4.037021e−02 | G protein−coupled receptor 183 |
| Syp | −0.2905242 | 4.081264e−02 | synaptophysin |
| Atp1a3 | −0.3693865 | 4.098119e−02 | ATPase, Na+/K+ transporting, alpha 3 polypeptide |
| Hap1 | −0.4010298 | 4.184648e−02 | huntingtin−associated protein 1 |
| Rin2 | −0.3161851 | 4.223955e−02 | Ras and Rab interactor 2 |
| Pagr1a | 0.8406428 | 4.320398e−02 | PAXIP1 associated glutamate rich protein 1A |
| 9930012K11Rik | −0.3244023 | 4.364790e−02 | RIKEN cDNA 9930012K11 gene |
| Fn1 | −0.3854521 | 4.459221e−02 | fibronectin 1 |
| Megf8 | −0.4036049 | 4.564958e−02 | multiple EGF−like−domains 8 |
| Exo1 | 0.4739798 | 4.571888e−02 | exonuclease 1 |
| Plekhg3 | −0.2936626 | 4.610656e−02 | pleckstrin homology domain containing, family G (w |
| B3glct | −0.2692229 | 4.714533e−02 | beta−3−glucosyltransferase |
| Zfp759 | −0.3113427 | 4.793228e−02 | zinc finger protein 759 |
| Apol10b | 0.4274514 | 4.829478e−02 | apolipoprotein L 10B |
| Pigg | 0.2844625 | 4.910487e−02 | phosphatidylinositol glycan anchor biosynthesis, c |

C

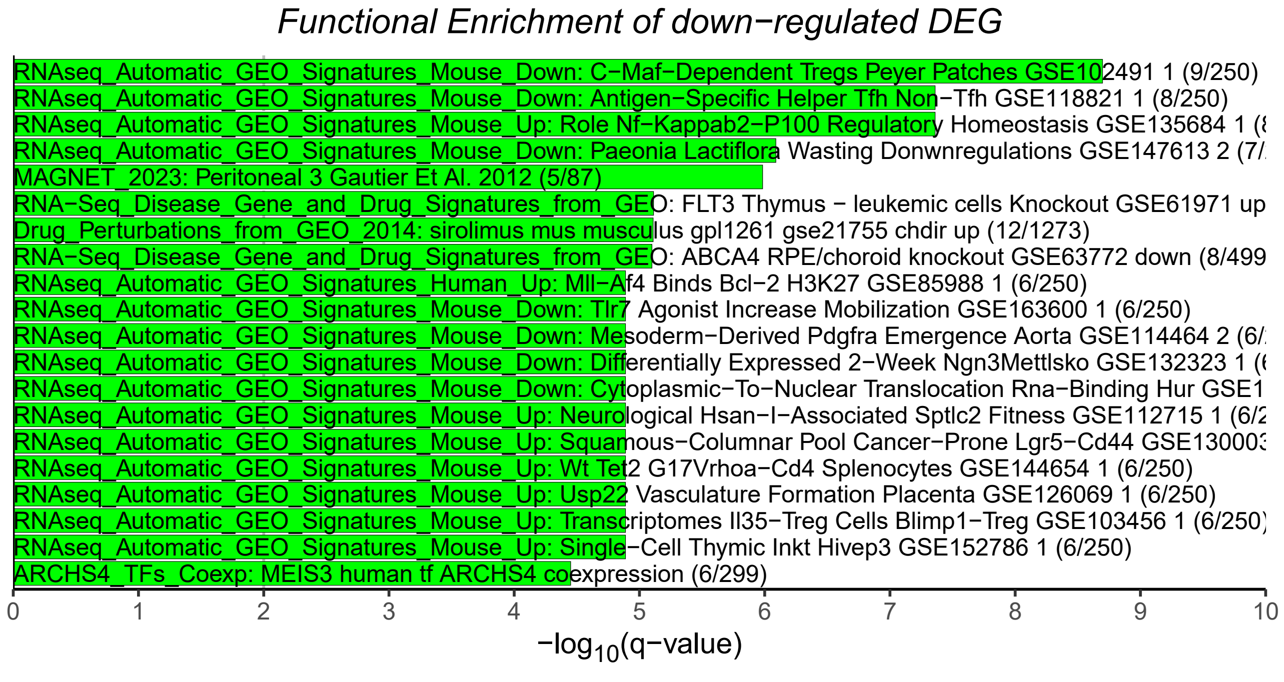


D

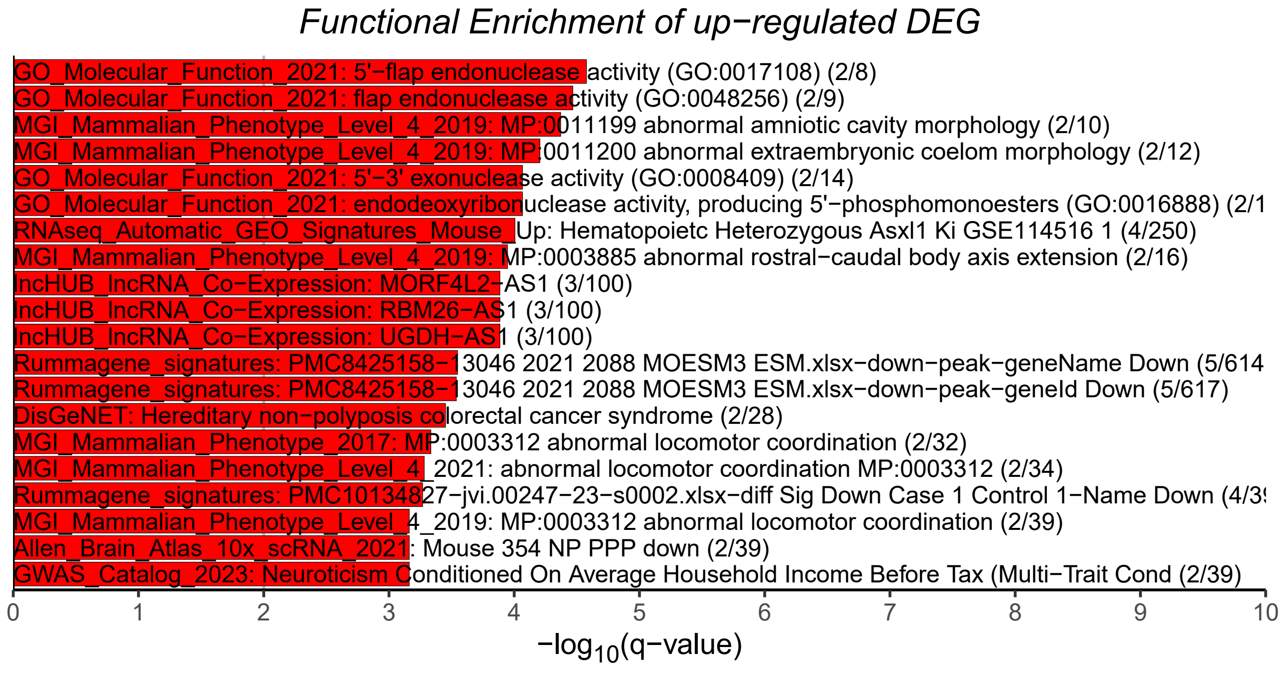

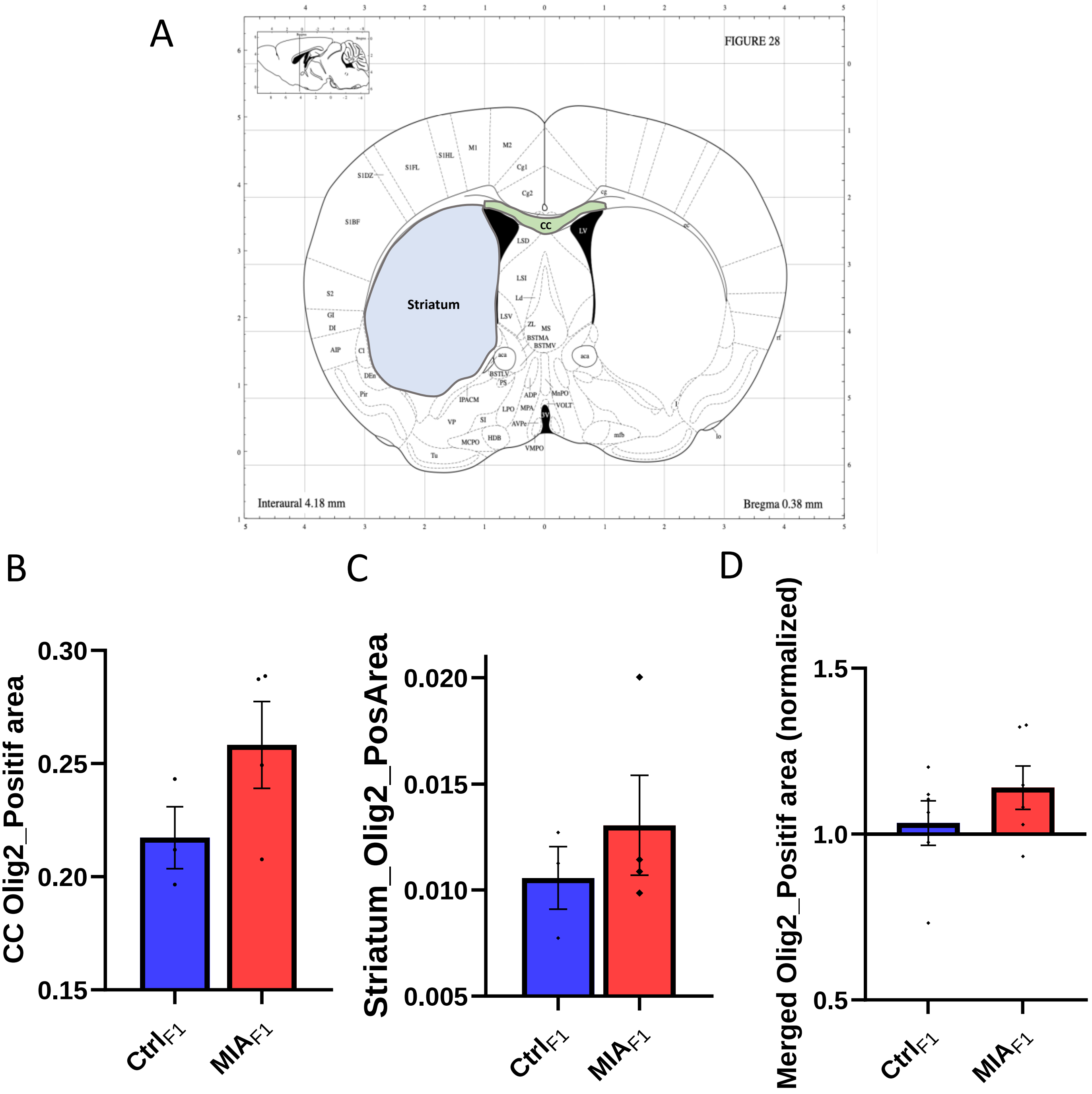

A
FIGURE 28
CC
LV
Striatum
Interaural 4.18 mm
Bregma 0.38 mm
B
CC Olig2_Positif area
0.30
0.25
0.20
0.15
Ctrl$_{F1}$
MIA$_{F1}$
C
Striatum_Olig2_PosArea
0.020
0.015
0.010
0.005
Ctrl$_{F1}$
MIA$_{F1}$
D
Merged Olig2_Positif area (normalized)
1.5
1.0
0.5
Ctrl$_{F1}$
MIA$_{F1}$

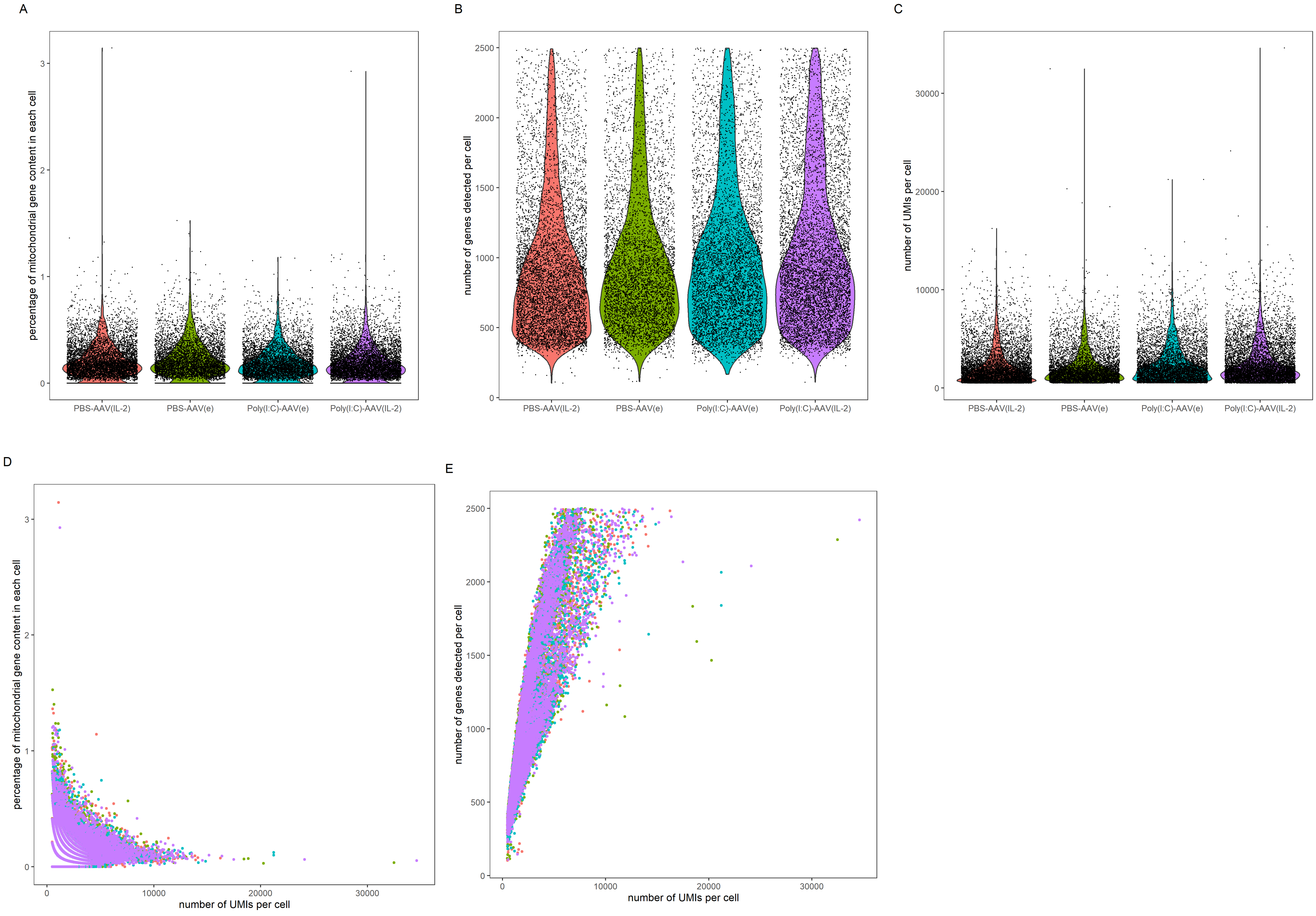

A
B
C
D
E
percentage of mitochondrial gene content in each cell
number of genes detected per cell
number of UMIs per cell
PBS-AAV(IL-2)
PBS-AAV(e)
Poly(I:C)-AAV(e)
Poly(I:C)-AAV(IL-2)

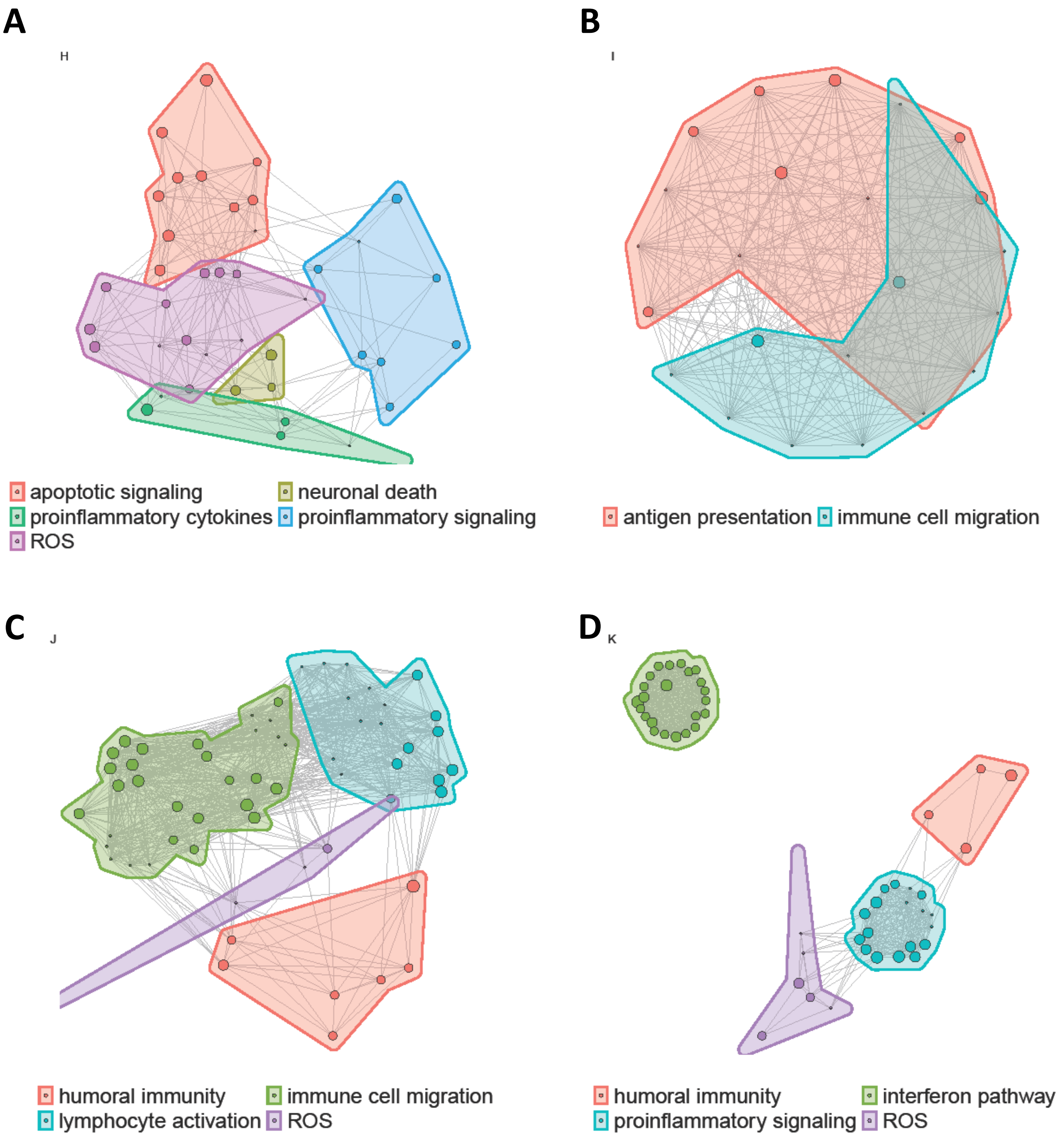
A
H
apoptotic signaling
neuronal death
proinflammatory cytokines
proinflammatory signaling
ROS
B
I
antigen presentation
immune cell migration
C
J
humoral immunity
immune cell migration
lymphocyte activation
ROS
D
K
humoral immunity
interferon pathway
proinflammatory signaling
ROS

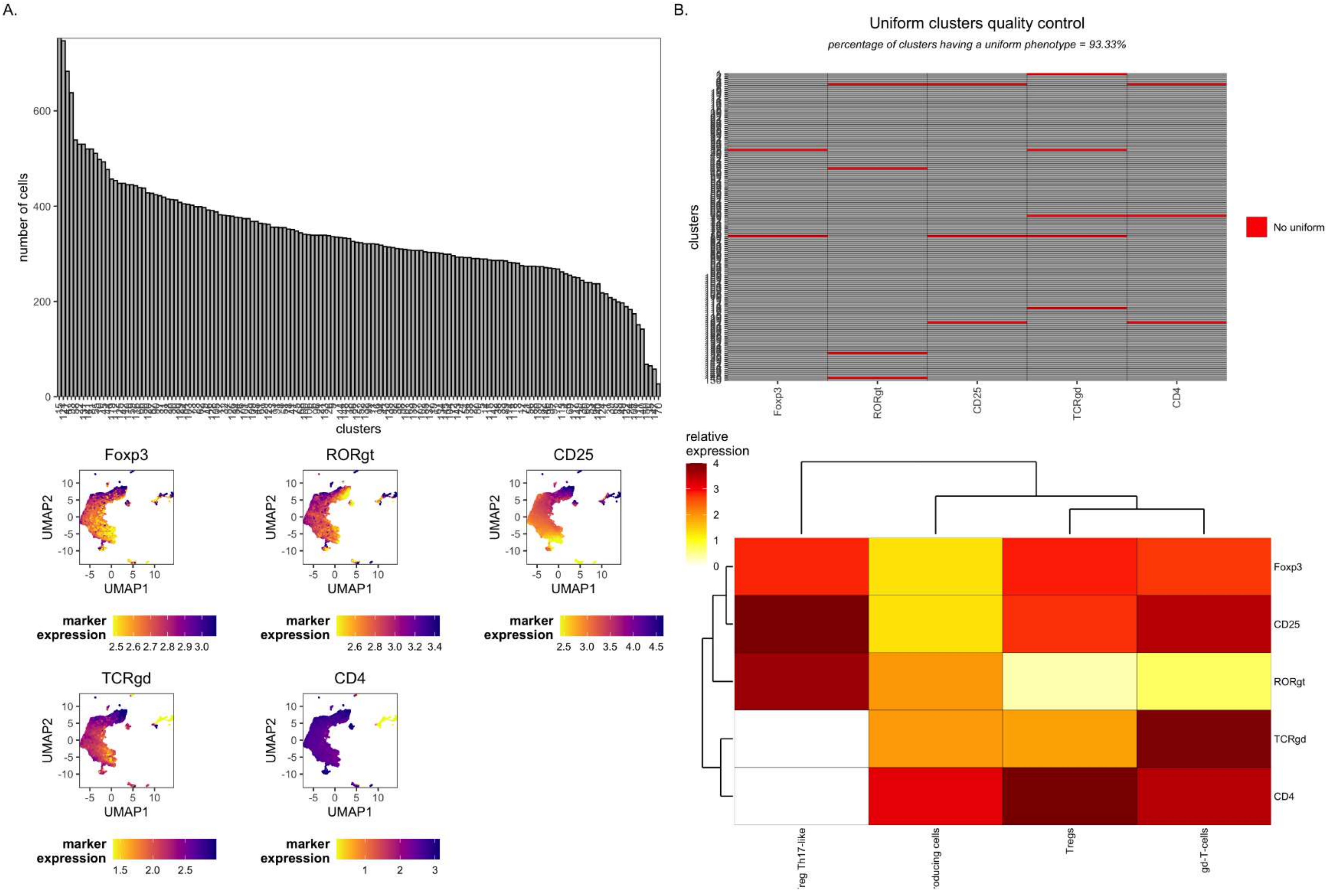

A.
B.
number of cells
clusters
Foxp3
RORgt
CD25
TCRgd
CD4
UMAP1
UMAP2
marker expression
Uniform clusters quality control
percentage of clusters having a uniform phenotype = 93.33%
No uniform
relative expression
Tregs
gd-T-cells

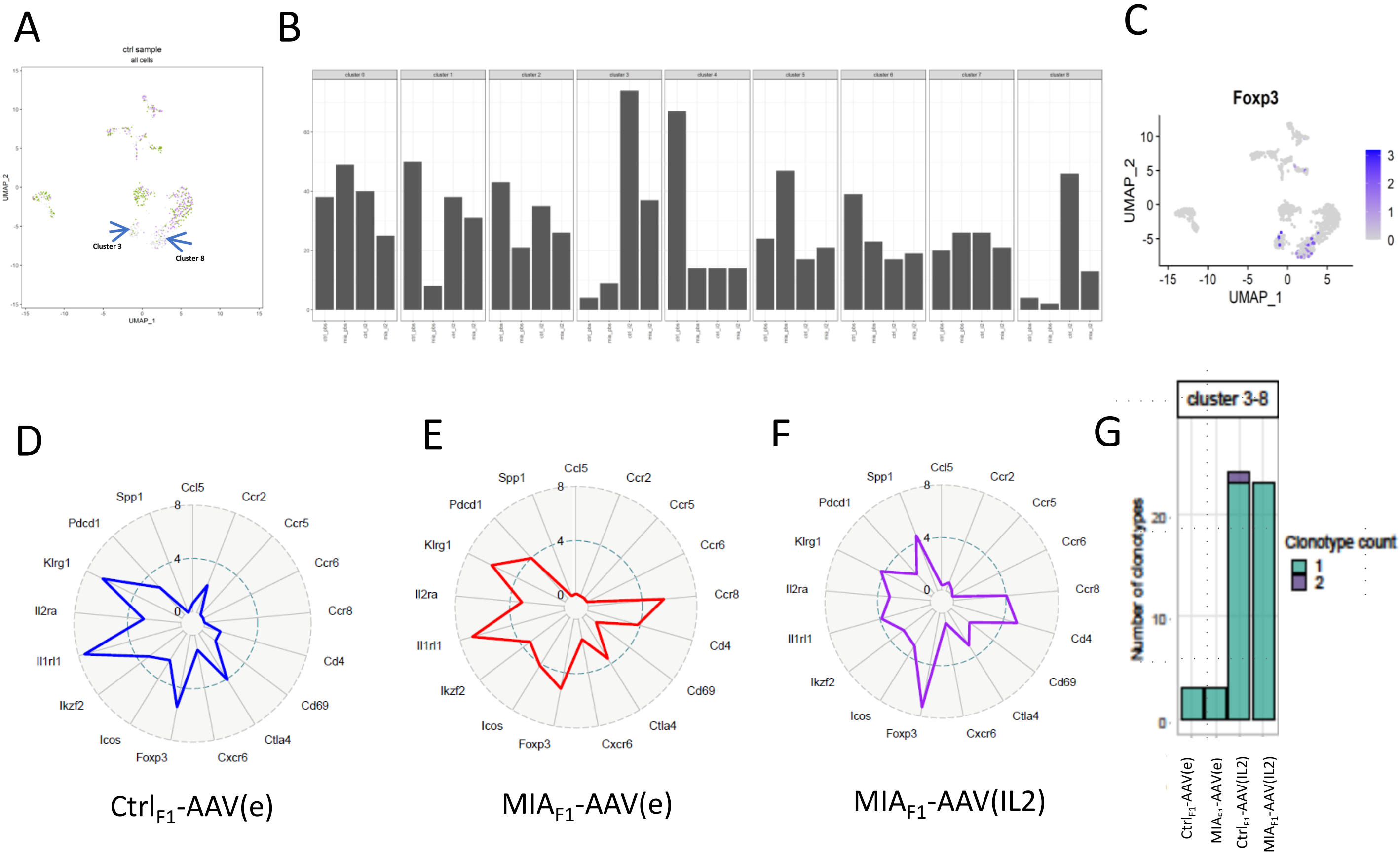
A
ctrl sample
all cells
UMAP_2
UMAP_1
Cluster 3
Cluster 8
B
C
Foxp3
UMAP_2
UMAP_1
D
Ccl5
Ccr2
Ccr5
Ccr6
Ccr8
Cd4
Cd69
Ctla4
Cxcr6
Foxp3
Icos
Ikzf2
Il1rl1
Il2ra
Klrg1
Pdcd1
Spp1
Ctrl$_{F1}$-AAV(e)
E
MIA$_{F1}$-AAV(e)
F
MIA$_{F1}$-AAV(IL2)
G
cluster 3-8
Number of clonotypes
Clonotype count
1
2
Ctrl$_{F1}$-AAV(e)
MIA$_{F1}$-AAV(e)
Ctrl$_{F1}$-AAV(IL2)
MIA$_{F1}$-AAV(IL2)

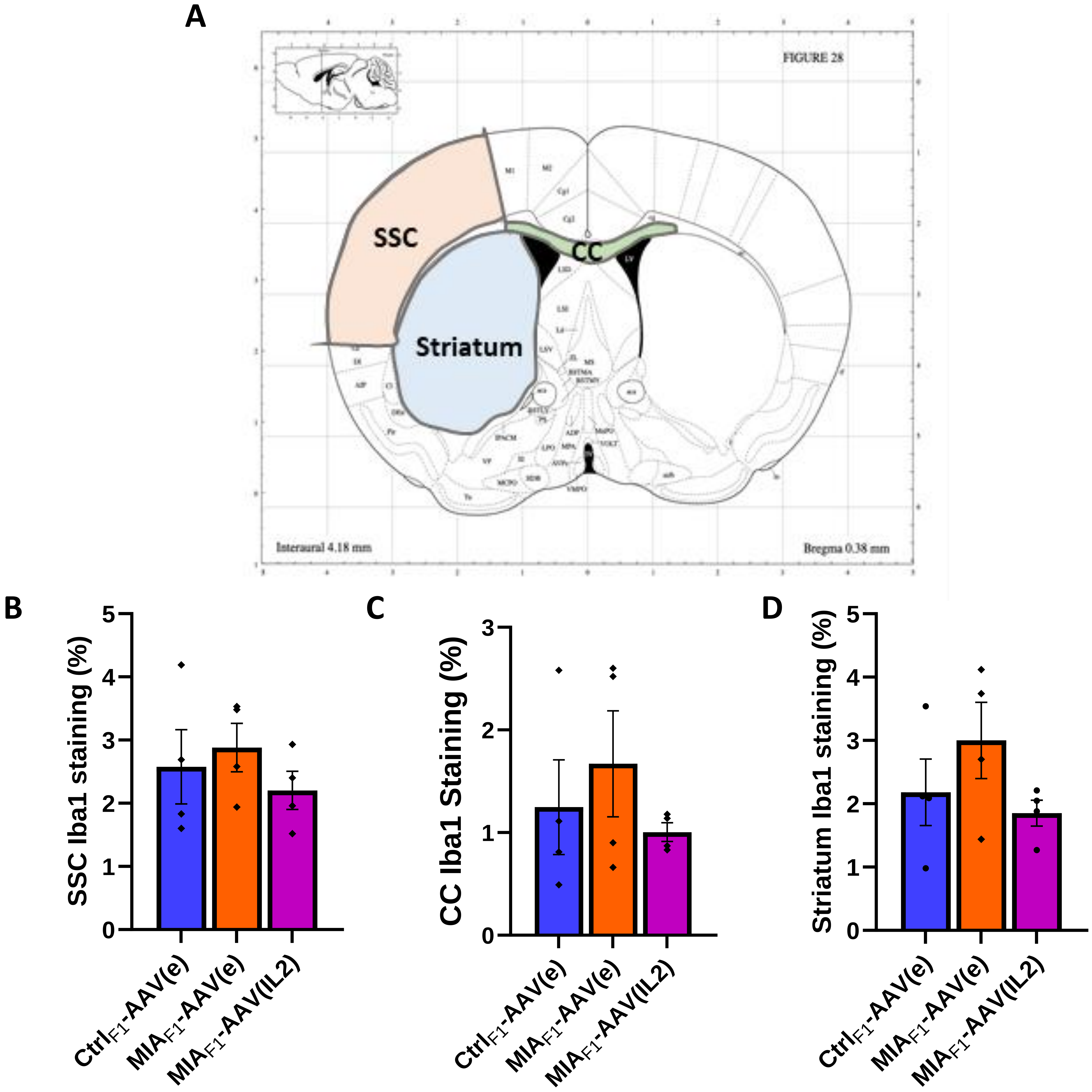
A
FIGURE 28
SSC
CC
Striatum
Interaural 4.18 mm
Bregma 0.38 mm
B
SSC Iba1 staining (%)
0
1
2
3
4
5
Ctrl$_{F1}$-AAV(e)
MIA$_{F1}$-AAV(e)
MIA$_{F1}$-AAV(IL2)
C
CC Iba1 Staining (%)
0
1
2
3
Ctrl$_{F1}$-AAV(e)
MIA$_{F1}$-AAV(e)
MIA$_{F1}$-AAV(IL2)
D
Striatum Iba1 staining (%)
0
1
2
3
4
5
Ctrl$_{F1}$-AAV(e)
MIA$_{F1}$-AAV(e)
MIA$_{F1}$-AAV(IL2)

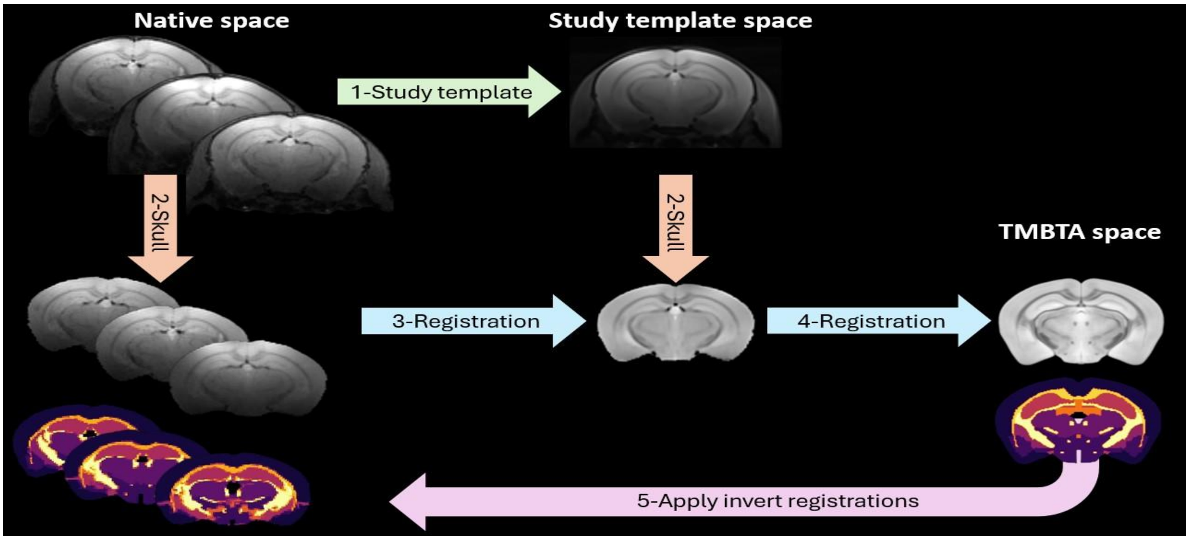
Native space
Study template space
1-Study template
2-Skull
2-Skull
TMBTA space
3-Registration
4-Registration
5-Apply invert registrations

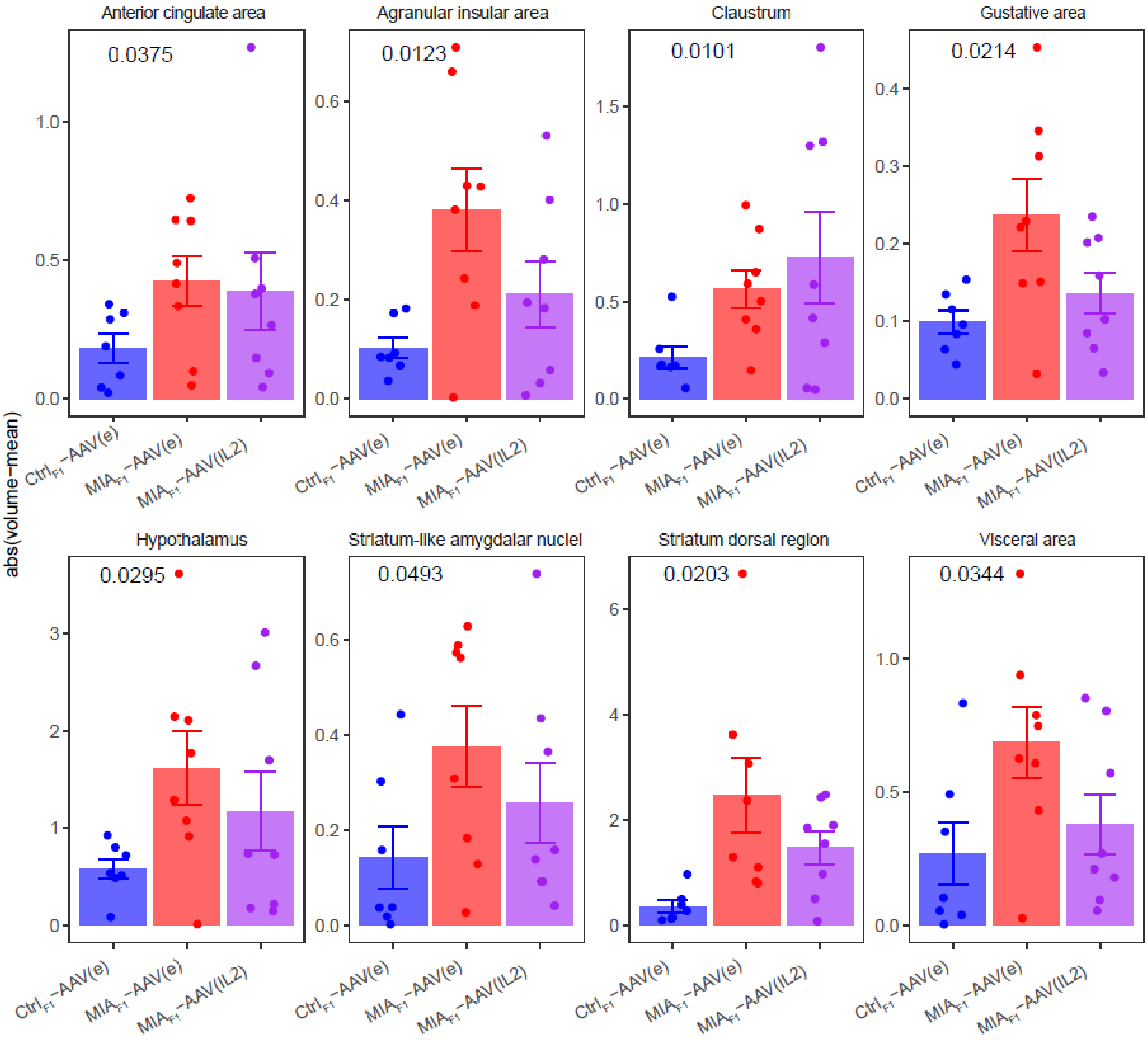
abs(volume−mean)
Anterior cingulate area
0.0375
Agranular insular area
0.0123
Claustrum
0.0101
Gustative area
0.0214
Hypothalamus
0.0295
Striatum-like amygdalar nuclei
0.0493
Striatum dorsal region
0.0203
Visceral area
0.0344
$Ctrl_{F1}$−AAV(e)
$MIA_{F1}$−AAV(e)
$MIA_{F1}$−AAV(IL2)
